# Ischemic and hemorrhagic stroke lesion environments differentially alter the glia repair potential of neural progenitor cell and immature astrocyte grafts

**DOI:** 10.1101/2023.08.14.553293

**Authors:** Honour O Adewumi, Gabriela I Berniac, Emily A McCarthy, Timothy M O’Shea

## Abstract

Using cell grafting to direct glia-based repair mechanisms in adult CNS injuries represents a potential therapeutic strategy for supporting functional neural parenchymal repair. However, glia repair directed by neural progenitor cell (NPC) grafts is dramatically altered by increasing lesion size, severity, and mode of injury. To address this, we studied the interplay between astrocyte differentiation and cell proliferation of NPC *in vitro* to generate proliferating immature astrocytes (ImA) using hysteretic conditioning. ImA maintain proliferation rates at comparable levels to NPC but showed robust immature astrocyte marker expression including Gfap and Vimentin. ImA demonstrated enhanced resistance to myofibroblast-like phenotypic transformations upon exposure to serum enriched environments *in vitro* compared to NPC and were more effective at scratch wound closure *in vitro* compared to quiescent astrocytes. Glia repair directed by ImA at acute ischemic striatal stroke lesions was equivalent to NPC but better than quiescent astrocyte grafts. While ischemic injury environments supported enhanced survival of grafts compared to healthy striatum, hemorrhagic lesions were hostile towards both NPC and ImA grafts leading to poor survival and ineffective modulation of natural wound repair processes. Our findings demonstrate that lesion environments, rather than transcriptional pre-graft states, determine the survival, cell-fate, and glia repair competency of cell grafts applied to acute CNS injuries.

## 1. Introduction

Traumatic injury to the adult central nervous system (CNS), such as following stroke or spinal cord injury (SCI), causes damage to neural tissue which does not spontaneously regenerate^1,2^. A biologically conserved multicellular wound repair response is initiated after CNS injury which protects adjacent spared neural tissue, clears damaged tissue debris, and generates a fibrotic scar that imparts structural and mechanical stability to the damaged tissue^1–11^. CNS injury lesions created by this natural wound repair response persist chronically, serve no neurological function, and do not support natural or stimulated plasticity^1,3,12^. For over 100 years there has been interest in grafting or transplanting neural cells at CNS injuries to restore lost neurological functions^13–17^. Despite progress over that time, we still lack sufficient mechanistic understanding in two key areas: (i) the graft phenotypes and functions necessary for supporting long-lasting neural regeneration after CNS injury, and (ii) the cell autonomous and non-autonomous regulation of cell graft functions in different CNS injury contexts^18^.

Successful axon regeneration strategies in the mammalian adult and insights from pro-regenerative organisms point towards the restoration of glia support networks as a promising goal for cell grafting. For example, studies manipulating intrinsic programs in spared neurons to stimulate axon regeneration have been most effective when there is sufficient preservation of glial cells to support the axon regrowth^19–22^, whereas there is limited regeneration when the same strategies are applied to large complete injury models where axon regrowth fails to traverse non-neural fibrotic lesion cores^23^. Propriospinal axon regeneration across complete SCI lesions can be achieved through fibrotic lesions by reinstating several developmentally essential molecular cues for axon growth back into the lesion core environment in combination with intrinsic neuronal growth stimulation^24^, but persistence of functioning circuits long-term requires repopulation of lesion tissue with glia for support^18^. Scar-free glia-based neural parenchymal repair that supports axon regeneration is observed in amphibians, zebrafish and notably in mammalian neonates^25^, which enables recovery of function that persists throughout adulthood^9,25–29^. Grafting cells acutely after injury to mimic neonate-like glia-based repair processes and provide the necessary glia support network represents a promising strategy for improving regeneration after adult CNS injury^18^. While glia progenitors or immature astrocytes have been grafted previously across various models of CNS injury, these studies have focused on replacing lost glia in preserved neural tissue or directing remyelination of regrowing axons which has meant grafting at later sub-acute timepoints^30–34^. Limited attention has been given to using these types of grafts as part of a strategy to modulate the natural wound repair responses in the acute setting of injury as a way of directing scar-free glia-based repair like that observed in the neonate. New methods for efficiently generating proliferating immature astroglia can help us to make progress in this area^18,35^.

There are many different sources of cells that have been explored for grafting in CNS injuries including fetal cells^36–41^, adult neural progenitor cells (NPC)^42–44^, and NPC derived from lines of embryonic stem cells (ESC) or induced pluripotent stem cells (iPSC)^45–50^. Irrespective of their source, cells are typically prepared for grafting via a period of *in vitro* culturing in serum-free environments with supportive supplements. Upon grafting into CNS injury lesions, grafted cells experience an abrupt environment change that can alter their phenotypes in significant ways. Recently, we derived mouse RiboTag NPC from mouse embryonic stem cells (mESC)^18,51^ and used these cells as a biological tool to begin dissecting how cell graft phenotypes are shaped by acute lesion environments^18^. The RiboTag system permits selective isolation of mRNA from grafted NPC *in vivo* for transcriptomic analysis and evaluation of the spatial heterogeneity of grafts by using RiboTag as an immunohistochemistry (IHC) reporter, thereby affording detailed insights into the phenotype and functions of grafted cells. Using RiboTag NPC we demonstrated that grafting into acute CNS injuries directs cells towards wound repair astroglial phenotypes by non-cell autonomous lesion cues to generate graft progeny that share morphological, transcriptional, and functional similarities with host astrocytes responding to CNS injury.

Grafting NPC into lesions at two days after injury contributed to reducing the size of ischemic stroke and spinal cord injury lesion cores^18^, but the effectiveness of the graft derived glia-based repair varied across these two CNS injury types, with moderate L-NIO induced ischemic stroke lesions being completely filled with graft-derived glia whereas severe SCI permitted only a few graft-derived glia to form bridges across lesions^18^. Although severe lesions appear to dysregulate glia repair by NPC grafts, how such environments alter lineage restricted glia progenitors or immature astrocytes has not yet been studied and identification of the lesion cues responsible for dysregulation of graft-derived glia repair remains elusive.

Here, we optimized a new methodology for deriving proliferating immature astrocytes (ImA) for grafting into CNS injuries by applying a hysteretic conditioning paradigm to NPC. We evaluated differences in grafting outcomes between these lineage-restricted ImA and multipotent NPC in ischemic and hemorrhagic striatal stroke injuries. Compared to NPC, ImA showed enhanced resistance to myofibroblast-like phenotypic transformations upon exposure to serum enriched environments *in vitro* and were more effective at wound closure *in vitro* compared to quiescent astrocytes (qA). ImA grafted into ischemic strokes directed effective glia-based repair comparable in nature to that observed for NPC but both graft candidates showed poor repair outcomes in more severe hemorrhagic injuries. Our findings demonstrate the feasibility of using hysteretic conditioning to generate immature astrocytes for CNS injury grafting and further refine our understanding of how different lesion environments instruct graft phenotypes and functions.

## 2. Materials and Methods

### 2.1. Culture of Neural Progenitor Cells (NPC)

Female mouse embryonic stems cells (mESC) that express the hemagglutinin (HA) epitope tag on modified ribosomal protein L22 (Rpl22) (RiboTag) were used to generate NPC by neural induction and expansion as previously described^18^. Briefly, mESCs were cultured on gelatin coated flasks (0.1% gelatin, Sigma, Lot# SLCK3991, solubilized in ultra purified water for 30 minutes at 50°C and sterile filtered) in mESC media containing KnockOut™ DMEM media (Cat# 10829018, ThermoFisher Scientific), 15% Fetal bovine serum (FBS) (ES cell tested) (ThermoFisher Sci Cat# 10439-016), MEM Non-Essential Amino Acids Solution (100X) (Cat# 11140050, ThermoFisher Scientific), Antibiotic-Antimycotic (100X) (Cat# 15240096, ThermoFisher Scientific), 100X EmbryoMax® Nucelosides (EMD Millipore Cat# ES-008-D), Beta-Mercaptoethanol (Sigma Aldrich M3148) and Leukemia inhibitory factor (LIF) (1 million units/mL stock) (EMD Millipore Cat# ESG1106). Embryoid bodies were formed by the removal of LIF and FBS supplements for 2 days followed by a four day period of culture in differentiation media which contained Advanced DMEM/F12 (ThermoFisher Sci Cat# 12634-010) supplemented with L-Glutamine (Thermofisher Scientific Cat# 25-030-081) (2mM) Knockout serum replacement (ThermoFisher Sci Cat# 10828010), retinoic acid (RA) (R2625-50MG, Sigma) (50nM) and purmorphamine (PUR) (sonic hedgehog (SHH) agonist) (SML0868-5MG, Sigma) (500nM). Neurally induced cells were expanded in Neural Expansion (NE) media comprised of Advanced DMEM/F12 supplemented with B27 (no Vitamin A) (50X) (ThermoFisher Sci Cat # 12587010), Antibiotic-Antimycotic (100X) (Cat# 15240096, ThermoFisher Scientific), Nucleosides (EmbryoMax, 100X Nuclesides Cat #ES-008), 100 ng/ml of Epidermal Growth Factor (EGF) (Peprotech, Cat# AF-100-15-100UG) and 100 ng/ml of Fibroblast Growth Factor (FGF) (Peprotech, Cat# 100-18B-100UG). NPC were passaged every 2-4 days by trypsinization in 0.05% trypsin EDTA (obtained from 0.25% Trypsin EDTA Gibco #25200056, diluted with 1X PBS (Gibco, #10010049)) for 3 minutes and centrifuged at 1000 rpm to collect the cell pellets. NPC used in this study were maintained at a passage number of 30 or less.

### 2.2. Differentiation and Hysteretic Conditioning of NPC

NPC are normally cultured in NE Media supplemented with 100 ng/ml of EGF and FGF (E/F). NPC were differentiated in a T-25, T-75 or T-150 culture flask at 140,000 cells per cm^2^ by culturing cells in Neural expansion media supplemented with either 1, 10, 25, or 50 ng/ml of E/F, 1% FBS (Gibco, REF 10437-010-100mL), 100 ng/ml of Ciliary Neurotrophic factor (CNTF) (Peprotech, Cat# 450-113-100UG), or combination of these morphogens for 2, 4, 6, or 8 days.

To determine cell growth rates, cells were counted at designated timepoints using a hemocytometer. Total cell number was obtained and the growth rate formula, 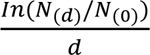 was utilized where *N(d)* is the number at the day of interest, *N(0)* is the starting number and *d* is the difference of days between the starting day and the day of interest.

For hysteretic conditioning of NPC, astrocyte differentiation was performed for 2 days by exposure to 1% FBS, CNTF, or 1%FBS +CNTF, cells were reintroduced into NE media supplemented with 100 ng/ml E/F for 3 days. For multiple cycles of hysteresis, the cells were again reintroduced into media containing astrocyte differentiation morphogens followed by NE media supplemented with 100 ng/ml E/F which was repeated for the designated number of cycles.

### 2.3. *In vitro* coverslips and immunocytochemistry (ICC)

For *in vitro* studies, 30,000 to 50,000 cells were seeded onto 10 mm round coverslips (Thorlabs) that were coated with 0.1% gelatin and placed into individual wells of a 24 well plate. Cells were cultured on gelatin coated coverslips for 2-4 days in 1 ml of media. At designated endpoints, the media was removed, the coverslips were rinsed with 1X PBS and then fixed with freshly prepared 4% paraformaldehyde (from 32% Paraformaldehyde Solution, Electron Microscopy Science Cat# 15714-S) for 30 minutes. For ICC, the coverslips were collected and rinsed 6 times with 1X Tris Buffered Saline (TBS) buffer, blocked for 1 hour in 5% Normal donkey serum (NDS, Jackson ImmunoResearch Laboratories, Lot# 161590) and permeabilized using 0.5% Triton x-100 in 1X TBS then incubated in primary antibody solution (with corresponding dilutions of each antibody in 1X TBS with 0.5% Triton x-100) overnight. The next day, the coverslips were washed 6 times, incubated in secondary antibody solution (1:250 dilution of each secondary antibody, 5% NDS, 5% 1X TBS with 0.5% Triton x-100 in 1X TBS) for 2 hours before then being washed again 6 times. The coverslips were then incubated in DAPI solution (2ng/ml; Molecular probes) for 10 minutes, rinsed 3 times and mounted onto microscope slides using Prolong Gold antifade reagent (Invitrogen, Cat# P36934) Slides were allowed to dry for at least 24 hours before being imaged using an Olympus IX83 Widefield Fluorescence microscope. Image analysis was carried out using FIJI/ImageJ.

The primary antibodies used in this study included: rat anti-Gfap (1:1000, Thermofisher, #13-0300); guinea pig anti-Gfap (1:1000, Synaptic Systems, #173-004); rabbit anti-Gfap (1:1000, Agilent #Z033401-2); rabbit anti-alpha-Sma (1:200, Thermofisher #PA5 16697); goat anti-HA (1:800, Novus, NB600-362); rabbit anti-HA (1:1000, Sigma, H6908-.2ML); rabbit-anti-Top2a (1:500, Abcam, ab52934); rat anti-Ki-67 (1:200, Invitrogen, 14-5698-82); goat anti-PDGFRa (1:500, R&D Systems, AF1062); guinea pig anti-Vimentin (1:500, Synaptic Systems, #172-004); goat anti-Nestin (1:500, R&D Systems, AF2736); rabbit anti-P2ry12 (1:800, Ana Spec, AS-55043A); goat anti-Cd13 (1:500, R&D Systems, AF2335); guinea pig anti-Iba-1 (1:500, Synaptic Systems, #234-004); goat anti-Sox9 (1:500, R&D Systems, AF3075); goat anti-Serpin A3N (1:200, R&D Systems, AF4709); rabbit anti-Id3 (1:200, Abcam, ab41834); guinea pig anti-NeuN (1:500, Synaptic Systems, #266-004); rat anti-P2ry12 (1:100, Biolegend, 848002); rat anti-Pu.1 (1:500, Biolegend, 681302); guinea pig anti-s100b (1:1000, Synaptic Systems, #287 004); rabbit anti-Atp1b2 (1:400, Abcam, ab185207); rabbit anti-Aldoc (1:400, Abcam, ab87122); goat anti-Clusterin (1:200, R&D Systems, AF797-SP); rat anti-Cd31 (MEC 13.3 clone) (1:200, BD Biosciences, 550274); sheep anti-S100A6 (1:200, R&D, AF4584). The secondary antibodies used were purchased from Jackson ImmunoResearch Laboratories with donkey host and target specified by the primary antibody. F-actin staining was done using Acti-stain 555 phalloidin (1:2000, Cytoskeleton, PHDH1A) and counter stained with DAPI.

### 2.4. RNA extraction and qPCR

Cell pellets (1-4 million cells total) were collected after trypsinization and centrifugation and stored at −80°C. RNA was extracted using the Qiagen RNeasy Plus Mini Kit (Cat# 74134). Briefly, cell pellets were resuspended in lysis buffer supplemented with beta-mercaptoethanol (Sigma, #63689) and homogenized using a handheld rotor with an attached pellet pestle for 30-45 seconds followed by a 2-minute high-speed centrifugation to remove insoluble cell debris.

The supernatant was then centrifuged through a gDNA eliminator spin column for 30 seconds and the flow through was mixed with 70% ethanol and passed again through an RNeasy column. The column was washed 3 times with wash buffer and dried for 2 minutes. The column was placed into a collection tube and centrifuged twice with RNase-free water to elute the purified RNA. All procedures were performed on ice and at a workbench space sprayed generously with RNase away. Extracted RNA was stored at −80°C and used within a month after extraction. The RNA concentration was measured using a nanodrop (Thermo Scientific Nanodrop 2000 spectrophotometer) with an RNA purity, determined by an 260nm/280nm ratio, of greater than 1.8 used for cDNA synthesis. To generate cDNA, RNA was diluted to a 500 ng/16 μl concentration using water and combined with 4 μl of iScript Reverse Transcription Supermix for RT-qPCR (Biorad, Cat# 1708841) and processed in a thermocycler (MJ Research PTC-200 Peltier Thermal Cycler) using manufacturer’s instructions. The cDNA was stored at −20°C until use. For qPCR of designated genes, primers were designed in house and purchased from Integrated DNA technologies; The NCBI ID for the gene of interest was identified and searched on the MGH Primer Bank for previously developed Primers (Supplementary Table 1). The selected forward and reverse primers were blasted on NCBI Blast to confirm gene specificity. The qPCR plate was set up such that each well contained 10 μl of total mixture; 5 μl of 2X SsoAdvanced Universal SYBR Green Supermix, 1 ul of Forward and Reverse primers for a 500 nM total concentration, 1 μl of cDNA and 2 μl of nuclease free water. Using the StepOnePlus RT-PCR System, samples were processed for 40 cycles at 95 °C for 30 seconds, 98 °C for 15 seconds and 60 °C for 60 seconds. Each sample was run in triplicates with the cycle threshold (ct) values normalized to the corresponding HA ct values as a housekeeping gene and then normalized to NPC to obtain the negative ΔΔ ct values for comparison.

### 2.5. Animals

All *in vivo* experiments and surgical procedures adhered to protocols approved by the Boston University IACUC (Protocol Numbers: PROTO202100013, PROTO202000045). Female C57BL/6 mice (JAX#000664) were received at 7 weeks of age and used for all *in vivo* experiments between 8-12 weeks of age. Animal Housing at the Boston University Animal Facility included a 12-hour light/dark cycle, controlled humidity, and temperature. Mice that underwent surgical procedures received analgesia pre- and post-surgery (buprenorphine, 0.1mg/kg). No animals in this study received immunosuppression drugs. All cell grafts were female cells that were transplanted into female mice to mitigate any sex-dependent effects that are established to alter graft outcome since sex mismatched cell grafts can provoke exacerbated macrophage, microglia and T cell responses^52^.

### 2.6. Surgery Procedures

Surgeries were performed under isoflurane (1.5-3%) and oxygen.

#### Craniotomy

A stereotaxic apparatus was used to stabilize the shaved mouse head. After sanitization with betadine and ethanol, an incision was made using stainless steel surgical blade and a 2 mm-by-2 mm craniotomy window was made over the left coronal suture using a surgical drill.

#### Stroke lesion models

Stroke lesions were made by injecting 1.5 μL of 27.4 or 40 mg/ml N5-(1-lminoethy)-L-ornithine (L-NIO) for ischemic strokes and 0.5 μL of 0.1 Units of Collagenase 1 solution (Gibco, Cat #17018-029) for hemorrhagic strokes at 0.15 μL/min. Injections were made to target the striatum at 1mm Anterior-Posterior (A/P), +2.5 mm Medial-Lateral (M/L) and 3 mm Dorsal-Ventral (D/V) from Bregma using a borosilicate pulled glass micropipette and a 10 μL Hamilton needle mounted to a syringe pump. Micropipettes were allowed to dwell in the brain for 4 minutes after the completion of the injection volume prior to slow removal from the brain using the stereotactic device. The skin was sutured with polypropylene sutures (UNIFT 6-0 #S-P618R13) after the injection and allowed to recover for 48 hours.

#### Cell grafting

NPC, ImA or qA were trypsinized and centrifuged to collect a cell pellet. 10 μl of 1 mg/ml E/F was added to each cell pellet before injection. Cells were mixed backloaded into pulled glass micropipettes. 1-1.2 μL (300,000 cells/μL) of cells were injected into healthy and injured mouse striatum at 0.15 μL/min. Injections were made to target the striatum at 1 mm A/P, +2.5 M/L and −3.5 D/V from Bregma. During the cell injections, the glass micropipette was moved dorsally 0.5mm at one-third volume of total volume (i.e 400 nl for 1.2 μL total volume) to ensure larger surface area distribution of the cell graft. The glass micropipette was allowed to rest in the brain for 4 minutes and slowly removed thereafter followed by skin closure by suturing as before.

#### Transcardial perfusions

At the designated experimental endpoint, isoflurane overdose was used to deeply anesthetize mice and cold PBS was transcardially perfused for 2 minutes followed by 7 minutes of perfusion with 4%PFA freshly made from 32% (Electron Microscopy Sciences Cat #15714-S) at a rate of 7 mL/min. Dissected brains were post-fixed in 4% PFA for 6-8 hours before being transferred into 30% sucrose solution and incubated at 4°C for 4 days with a sucrose solution change after 24 hours.

### 2.7. Histology and immunohistochemistry (IHC)

Brain tissue was cut coronally into 40 µm sections using a MICROM HM 525 or Leica CM1950 Cryostat. Tissue sections were stored in 96 well plates immersed with 1X TBS and 0.001 % sodium azide solution at 4°C. For IHC, tissue sections were stained via a free-floating staining method outlined in detail previously^4,18^. Briefly, tissue sections underwent antigen retrieval in 1N HCl, were washed in 1X TBS three times and then blocked and permeabilized for one hour in 1X TBS containing 5% Normal donkey serum (NDS, Jackson ImmunoResearch Laboratories, Lot #161590) and 0.5% Triton x-100. Primary and secondary antibodies used were the same as listed above. Nuclei were stained with DAPI (4’6’-diamidino-2-phenylindole dihydrochloride.

ProLong Gold (Invitrogen, Cat# P36934) anti-fade reagent was used to mount the brain sections onto slides. The brain sections were imaged using an IX83 Olympus Microscope and processed using Cell Sens software. Detailed Orthogonal Projections were processed through Constrained Iterative Deconvolution using Cells Sens Software and assembled using ZEN Lite (Zeiss).

### 2.8. Scratch Assay

NPC, ImA or qA were seeded into 6 well plates at a density of 7×10^5^ cells per well. After 24 hours in normal media conditions for each cell type, a 200 μL plastic pipette tip was used to scratch horizontally through the well. Cells were then washed, and media changed to include 1%FBS and or E/F. Bright field images of cell migration responses were collected immediately after scratch and at designated time points up to 72 hours. Images were appropriately thresholded and measured to determine the cell filled area using FIJI/ImageJ software.

### 2.9. Quantitative analysis of ICC and IHC

All quantitative analysis of ICC and IHC images were performed using FIJI/ImageJ. ICC and IHC images were quantified by manual counts of cells, intensity quantification, as well as area under curve calculations on radial angle profile intensity traces using protocols outlined in detail previously^4,18^. Staining colocalization was assessed using the RG2B colocalization plug-in as described before ^4,18^.

#### 2.1.0. Statistical Analysis

Graphs were constructed using GraphPad prism. Statistical evaluations were performed by running Student’s t-test as well as One-way or Two-way ANOVA with Tukey post hoc analysis where appropriate. Significance for statistically tested variables was defined by a p-value less than 0.05. All graphs presented display mean and standard error of mean unless otherwise specified. Individual data points are overlaid across all graphs.

## 3. Results

### 3.1. Ischemic and hemorrhagic striatal stroke lesion environments differentially alter NPC graft outcome

We have previously noted differences between NPC grafting outcomes for ischemic stroke lesions that have limited bleeding and crush spinal cord injury (SCI) that have extensive bleeding and hemorrhagic necrosis^18^. NPC grafted into ischemic strokes readily fill the entire lesion core with Gfap-positive astroglia whereas in SCI there are only small numbers of these graft derived astroglia^18^. To test whether such altered grafting outcomes are caused primarily by lesion type differences and rule out any possible influence of anatomy (i.e., spinal cord vs brain) and injury mode (i.e., solution injection vs forcep crush), we grafted the same NPC line into either ischemic or hemorrhagic stroke injuries that were made in the same neuroanatomical location in the mouse striatum by stereotactic injection of different stroke inducing agents. Cells were grafted at 48 hours after stroke and grafting outcome evaluated by IHC at 2 weeks (Figure 1A). To induce ischemic injuries with limited bleeding, we injected the vasoconstrictive agent, L-NIO (1.5 µL) as we have done before^18^ and used two different concentrations (27.4 mg/ml and 40 mg/ml) of L-NIO to create moderate and severe ischemic injuries respectively. A collagenase solution (0.5-1 µL at 0.1 U/µL) which acts to enzymatically degrade structural matrix at blood vessels causing vessel rupture, was used to generate intracerebral hemorrhagic stroke injuries with extensive bleeding^53,54^. Stroke lesions made using these two methods are reliable in location and size but show different pathophysiology and lesion phenotypes (Figure 1B)^4,53–56^. Evaluating untreated strokes at different time points up to 2 weeks post injury revealed that hemorrhagic lesions had narrower and less dense astrocyte borders at the lesion edge, fewer persisting P2ry12- and Iba-1-positive microglia at borders, slower but greater recruitment of Cd13-positive peripheral immune cells to infarct regions, less extensive fibrotic scar formation, and greater contraction of striatal neural tissue compared to ischemic lesions (Figure 1B, S1A, 1SB). Evaluation of stroke lesions at earlier post-injury timepoints (2-7 days) also revealed notable differences in Cd31-positive angiogenesis between the two stroke types. Ischemic strokes evaluated at 5 days showed extensive angiogenesis in lesion cores, but hemorrhagic injuries maintained coagulated hematomas with only sparse immune cell infiltrates and limited neocapillaries detected up to at least 1 week post injury (Figure S1C). Vascular remodeling differences between the injury types were maintained up to 2 weeks post injury with the morphology of Cd31-positive angiogenic vessels in hemorrhagic injuries having a more hyper-intense and diffuse distribution compared with ischemic injuries (Figure S1A, 1SB).

**Figure 1.**
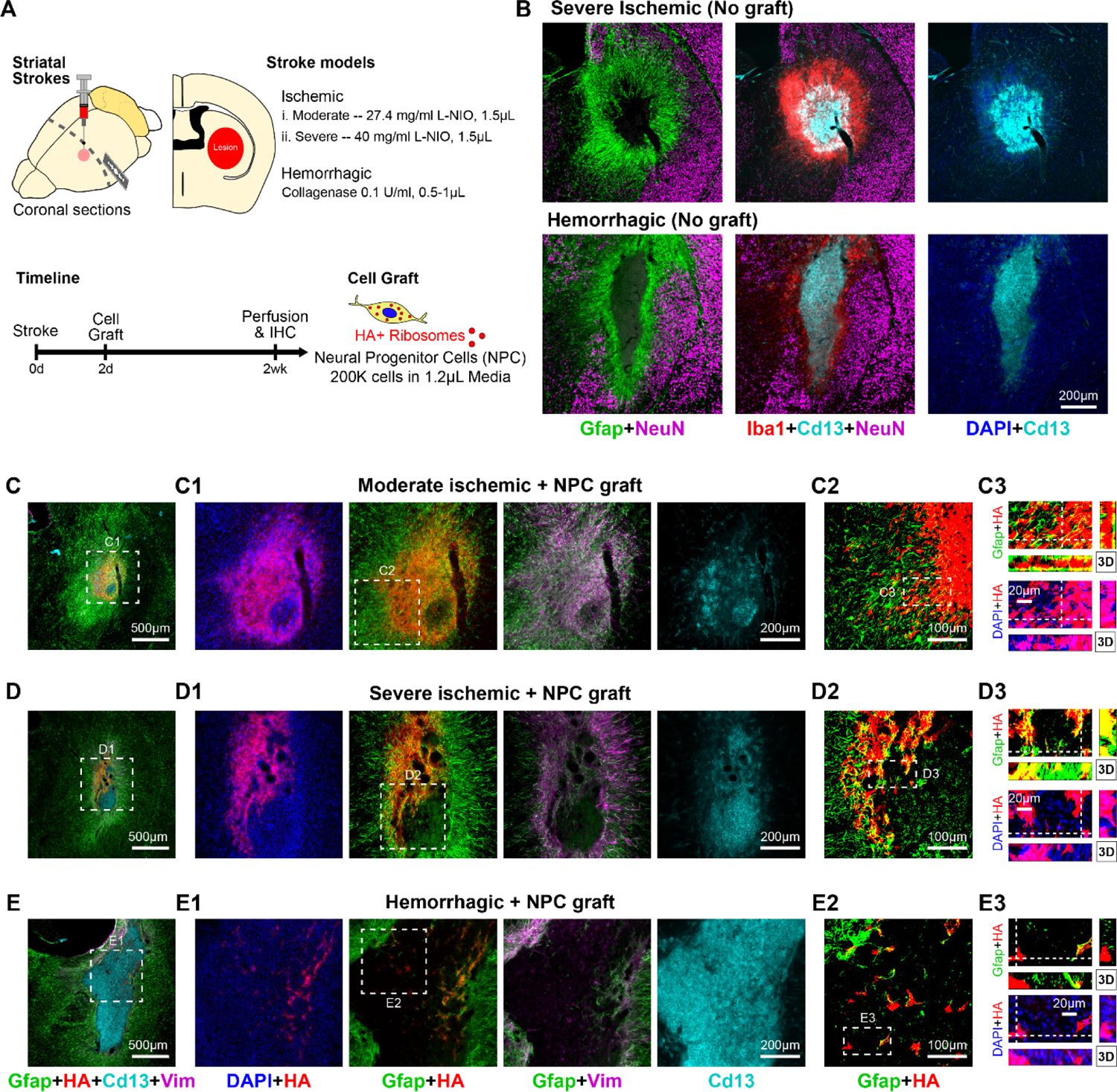
Ischemic and hemorrhagic striatal stroke lesion environments differentially alter NPC graft outcome. **A.** Schematic summarizing experimental approach for evaluating acute Ribo Tag NPC grafting outcomes in ischemic and hemorrhagic striatal strokes induced by injection of L-NIO and Collagenase solutions respectively. **B.** Survey immunohistochemistry (IHC) images of stroke lesions at 2 weeks showing differences in the astrocyte (Gfap), microglia (Iba1), and myeloid lineage cell (Cd13) responses as well as in the extent of contraction of NeuN-positive striatal tissue at ischemic and hemorrhagic lesions. **C-E.** Survey and detailed orthogonal IHC images showing NPC grafting outcome in moderate (**C**) and severe (**D**) ischemic strokes and hemorrhagic strokes (**E**). The severity and the type of stroke alters HA-positive NPC survival and wound repair astroglia phenotype (Gfap and Vimentin (Vim)) as well as the effectiveness of glia repair in Cd13-positive lesion cores.

Grafting NPC into the different stroke lesions showed that both size and mode of injury altered graft-derived glia-based repair outcomes (Figure 1C-E). NPC grafted into ischemic injuries were filled with NPC graft derived Gfap-positive astroglia that effectively reduced the numbers of Cd13-positive immune cells (Figure 1C). Increasing the severity of ischemic strokes resulted in less effective glia-based repair leading to pockets of Cd13-positive immune cells that were corralled by, but not infiltrated with, NPC graft progeny (Figure 1D). Compared to grafts made in either of the two types of ischemic injuries, NPC grafts in hemorrhagic strokes had significantly reduced cell survival, less effective glia-based repair, and numerous cells that were Gfap-negative suggesting that these cells did not differentiate into wound repair astroglia (Figure 1E). In hemorrhagic stroke lesions, the small numbers of surviving graft progeny persisted mostly at neural tissue margins of the lesion and were ineffective at corralling Cd13-positive immune cells that infiltrated in large numbers into lesion cores (Figure 1E).

These data show that glia-based repair outcomes directed by NPC grafts are altered by increasing lesion size, severity, and the different modes of stroke injury. Hemorrhagic lesion environments exceedingly perturb graft survival and wound repair astroglia functions compared to ischemic injuries. These data reinforce the notion that non-cell autonomous cues dominantly modulate NPC graft outcome and that strategies are needed to prime glia-based repair functions in cell grafts to overcome these lesion cues. To address this, we next focused on developing methods to generate immature astrocytes for grafting as we hypothesized that such cells may promote improved glia-based repair outcomes in severe CNS injuries where NPC are ineffective. The approaches taken to identify the methodology for deriving proliferating immature astrocytes as well as the outcomes of grafting immature astrocytes into stroke injuries compared to NPC are examined in more detail below.

### 3.2. Spontaneous glial differentiation of NPC *in vitro* requires concurrent suppression of proliferation programs

Early postnatal immature astroglia contribute to effective glia-based repair at CNS injury lesions^9,25–29^. These immature astroglia express many canonical astrocyte genes but, unlike adult astrocytes, also maintain a high proliferative state which likely contributes to their capacity to mount effective glia-based repair^57^. As a first approach to derive immature astroglia from NPC, we hypothesized that we could generate proliferating immature astroglia through small reductions in the concentration of EGF/FGF (E/F) mitogens applied to NPC cultures. Growth media that incorporates high concentrations of E/F (100 ng/ml) is required to keep ESC-derived NPC in their neural progenitor state and maintain high rates of proliferation *in vitro*^18^. Reducing the concentration of these mitogens 100-fold *in vitro* to 1 ng/ml evokes spontaneous, cell autonomously regulated differentiation of NPC over 4 days leading to a mixed glia culture of astrocytes and oligodendroglia (we have previously referred to this as SPONT) ^18^. However, the effect of intermediate E/F concentrations on glial cell differentiation or proliferation has yet to be evaluated. To probe these effects, we cultured NPC in media containing decreasing concentrations of E/F from 100 ng/ml to 1 ng/ml for 2 or 4 days and examined proliferation state (Figure 2) and glial cell differentiation (Figure 3). Exposing cells to mild trypsinization and counting the recovered cells revealed a significant decrease in cell proliferation capacity with reduced E/F concentrations (Figure 2A-C). E/F concentrations down to 25 ng/ml enabled cells to continue to proliferate, albeit at rates that were less than half of those observed under the standard NPC conditions (100 ng/ml E/F), resulting in greater numbers of recovered cells at 4 days compared to initial seeding. For 10 ng/ml E/F treatment, the total number of recovered cells remained the same as the initial seeding number, although there was an observable increase in cell number from 2 to 4 days suggesting some, but low, levels of proliferation under these conditions. For 1 ng/ml E/F treatment, cell numbers decreased dramatically by 92.5% of the initial seeding density at 2 days and remained essentially unchanged (93.3%) at 4 days (Figure 2C) suggesting cells had become mitotically inactive with this low concentration of E/F. The reduced cell proliferation associated with lower E/F concentrations coincided with significantly decreased protein and gene expression of proliferation markers Top2a (*Top2a*) and Ki67 (*Mki67*) as well as decreased gene expression of the canonical neural stem cell gene *Igfbp2* (Figure 2B, D-E). While over 88.9% of NPC under standard 100 ng/ml E/F culture conditions were Top2a-positive, less than 52.6% of cells were Top2a-positive at 10 ng/ml, and 2.5% of cells remained Top2a-positive at 1 ng/ml E/F. A similar E/F concentration dependency of expression was also observed for the Ki67 proliferation marker demonstrating complete loss of mitotic activity with E/F withdrawal (Figure 2B).

**Figure 2.**
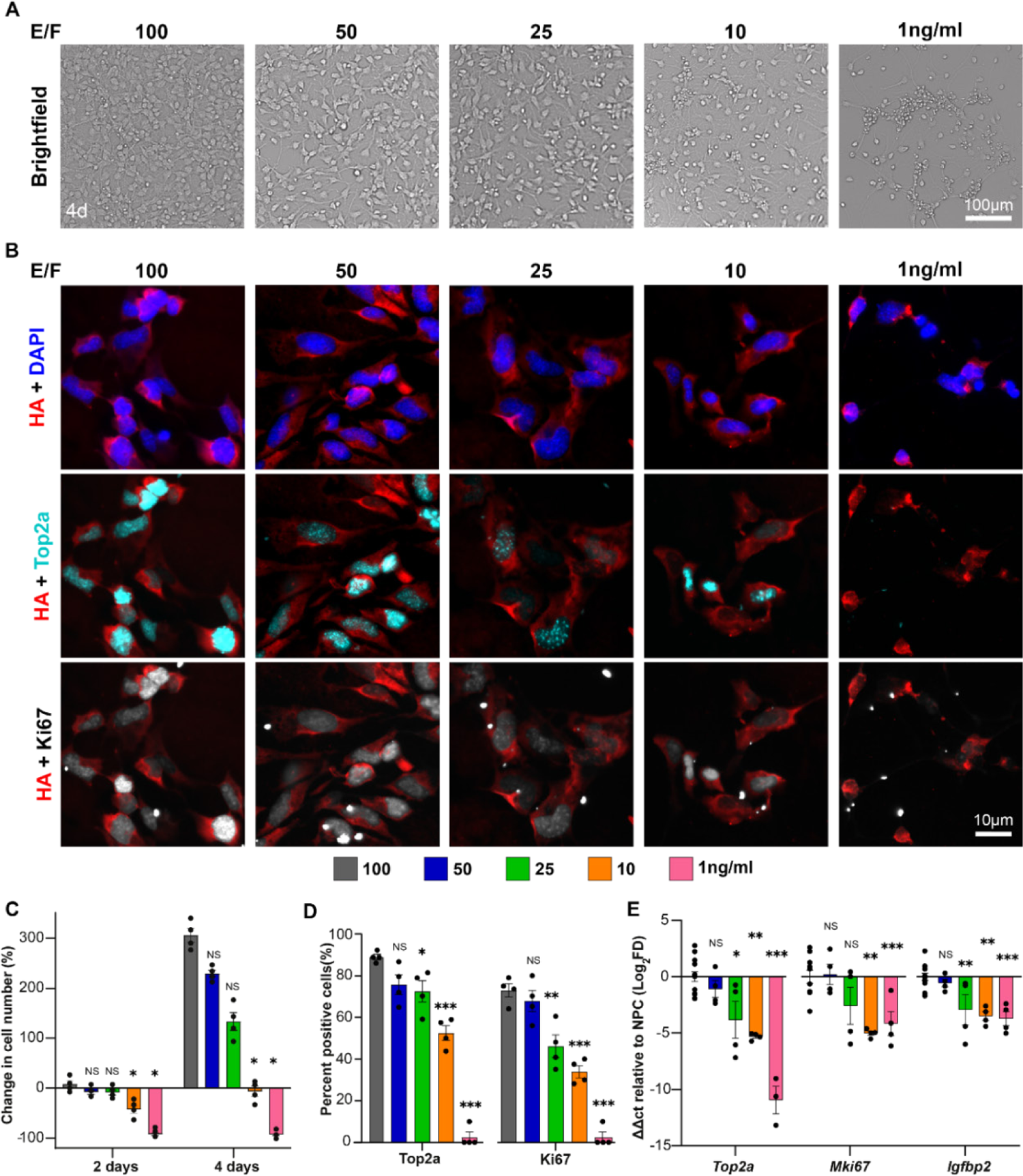
NPC proliferation state is directly dependent on EGF/FGF (E/F) concentrations. **A.** Brightfield microscopy images of NPC exposed to decreasing concentrations of E/F showing changes to cell density and morphology. **B.** ICC detailed images of E/F concentration-dependent changes in proliferation marker expression (Top2a and Ki67) in HA-positive Ribo Tag NPC. **C.** Percent change in cell number for decreasing concentrations of E/F at 2 and 4 days after seeding. Two-way ANOVA with Tukey multiple comparison test, not significant (NS), *p-value < 0.0005. Individual data points showing n=4 biological replicates per group. Sample are compared to NPC at 100 ng/ml (Grey). **D.** Quantification of the proportion of cells expressing proliferation markers (Top2a and Ki67) for different E/F concentrations. One-way ANOVA with Tukey test, not significant (NS), * p-value < 0.05, ** p-value < 0.002, *** p-value < 0.0001. Individual data point showing n=4 for all groups. Sample are compared to NPC at 100 ng/ml (Grey). **E.** qPCR of proliferation (*Top2a* and *Mki67*) and immaturity (*Igfbp2*) genes. ΔΔct values normalized to NPC (100ng/ml E/F) with *Rpl22-HA* used as a housekeeping gene. One-way ANOVA with Tukey test, not significant (NS), * p-value < 0.02, ** p-value < 0.009, *** p-value < 0.0008. Individual data point showing n=13 for NPC and n=4 for all other groups. Sample are compared to NPC at 100 ng/ml (Grey). All graphs are mean±s.e.m.

**Figure 3.**
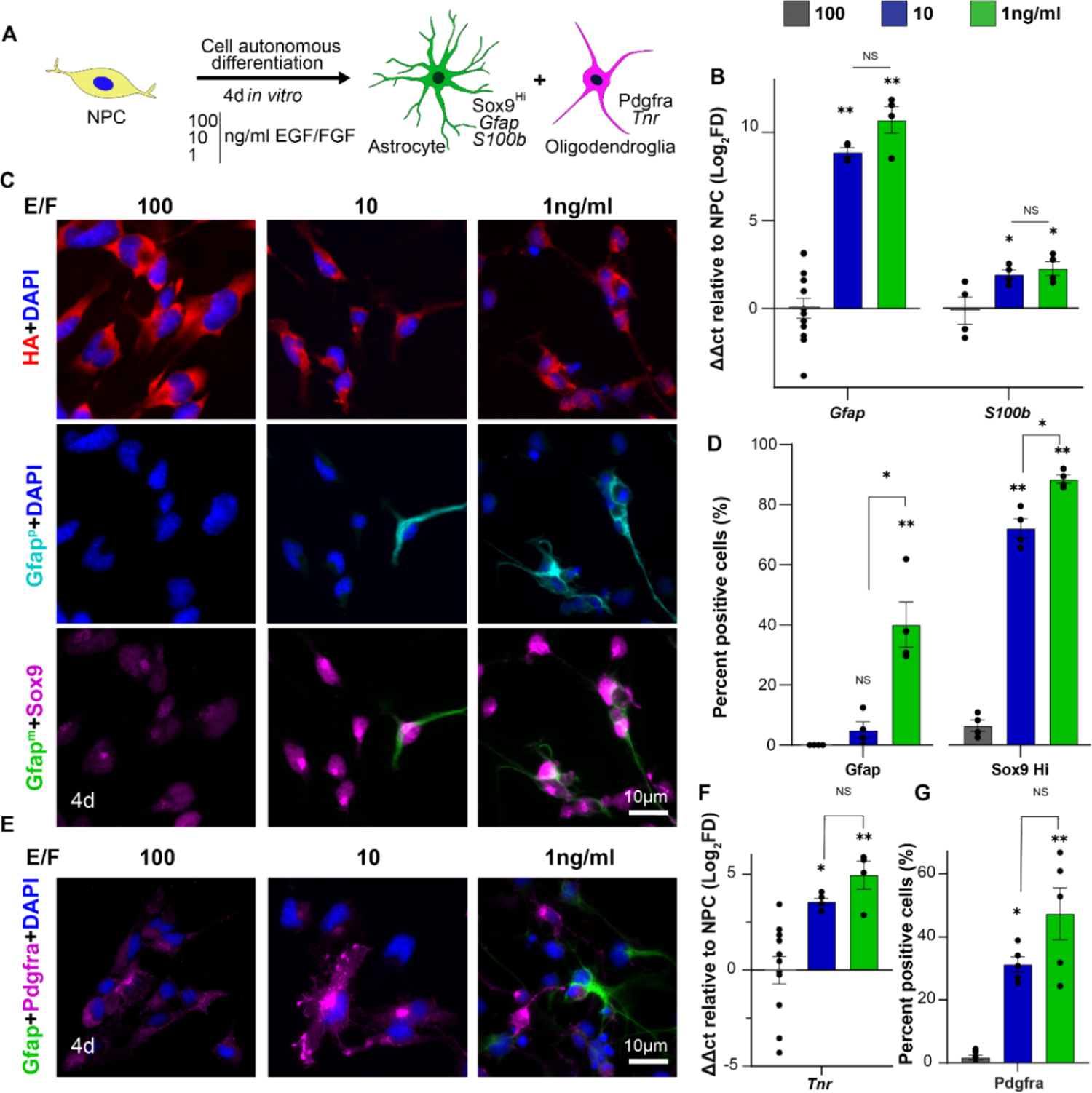
NPC differentiate into mixed glia cultures of astrocytes and oligodendroglia but only at low EGF/FGF (E/F) concentrations. **A.** Schematic of cell autonomous differentiation experimental approach. **B.** E/F concentration dependent changes in expression of astroglia (*Gfap* and *S100b*) genes by qPCR. ΔΔct values normalized to NPC (100 ng/ml E/F) with *Rpl22-HA* used as a housekeeping gene. One-way ANOVA with Tukey test, not significant (NS), * p-value < 0.05, ** p-value < 0.0001. Individual data point showing n=13 for *Gfap* 100ng/ml, n=3 for Gfap 10ng/ml, and n=4 for all other groups. Sample are compared to NPC at 100 ng/ml (Grey). **C.** ICC detailed images of E/F concentration-dependent differences in glial differentiation using polyclonal (Gfap^p^) and monoclonal (Gfap^m^) Gfap antibodies and Sox9. **D.** Quantification of the proportion of cells expressing astrocyte markers (Gfap and Sox9Hi). One-way ANOVA with Tukey test, not significant (NS), * p-value < 0.002, ** p-value < 0.004. Individual data point showing n=4 for all groups. Sample are compared to NPC at 100 ng/ml (Grey). **E.** ICC detailed images of E/F concentration-dependent differences in astrocyte (Gfap) and oligodendrocyte (Pdgfra) differentiation. **F.** E/F concentration dependent changes in oligodendroglia gene (*Tnr*) expression. ΔΔct values normalized to NPC (100 ng/ml E/F) with *Rpl22-HA* used as a housekeeping gene. One-way ANOVA with Tukey test, not significant (NS), * p-value < 0.02, ** p-value < 0.002. Individual data point showing n=13 for NPC and n=4 for all other groups. Sample are compared to NPC at 100 ng/ml (Grey). **G.** Quantification of the proportion of cells expressing oligodendroglia marker Pdgfra. One-way ANOVA with Tukey test, not significant (ns), *P < 0.003, **P < 0.0001. Individual data point showing n=4 for all groups. All graphs are mean±s.e.m. Sample are compared to NPC at 100 ng/ml (Grey).

To evaluate whether glial cell differentiation comparable to the SPONT condition (1 ng/ml E/F) can be achieved for any of the E/F concentrations that maintain proliferation capacity, we probed gene and protein expression for a variety of canonical astrocyte and oligodendrocyte markers (Figure 3). Astrocyte and oligodendroglia lineage maturation of NPC under SPONT differentiation conditions show characteristically high expression of *Gfap* and *Tnr* genes respectively, as identified by our previous single nuclei RNA-Seq experiments^18^, so transcript changes in these genes were used to assess glial cell differentiation for the five E/F concentrations. For both *Gfap* and *Tnr* genes, expression levels were inversely proportional to E/F concentrations (Figure 3B-F, S2E). Reducing E/F concentration by half to 50 ng/ml evoked a near 25-fold increase in *Gfap* gene expression compared to NPC but this was significantly lower than the 1675-fold increase in *Gfap* expression detected for the SPONT condition. Additional astrocyte enriched genes including *Emp3 and S100b* were also incrementally elevated by reducing E/F concentrations (Figure 3B, S2E). While reduced E/F concentrations resulted in measurable increased expression of these astrocyte genes, detectable protein expression of Gfap by immunocytochemistry (ICC) was only apparent in E/F concentrations of 10ng/ml or less (Figure 3C, D). Approximately 5% of cells at 10 ng/ml E/F and 40% of cells at 1 ng/ml E/F were Gfap-positive by ICC with both conditions showing detectable, but very low, levels of Gfap expression in positive cells (Figure 3C, D). The number and intensity of Gfap-positive cells detected in these conditions was consistent when probed with either polyclonal (p) or monoclonal (m) Gfap antibodies (Figure 3C). Other canonical healthy astrocyte markers such as S100b, Aldoc, Clusterin, and Atp1b2 were only detected in cells exposed to 1 ng/ml E/F condition (Figure S2A-C). Cells treated with low E/F concentrations showed increased expression of astrocyte nuclear marker, Sox9, above NPC baseline levels such that 72% of cells expressed Sox9 at high levels (Sox9^Hi^) with 10ng/ml EGF/FGF while 88.5% of cells were Sox9^Hi^ under 1ng/ml EGF/FGF conditions (Figure 3D). Reduced concentrations of E/F resulted in elevated *Tnr* gene levels and increased prevalence of Pdgfra-positive oligodendroglia cells such that over 47.3% of cells were Pdgfra-positive at 1 ng/ml (Figure 2F, G). Cells that were positive for glial cell markers by ICC under low E/F conditions did not co-express Ki67 (Figure S2D).

These data show that E/F mitogens added into NPC culture media have a concentration dependent effect on maintaining cell proliferation programs and that preserving proliferation capacity prevents glial differentiation of NPC *in vitro*. These data also show that glial differentiation of NPC *in vitro* requires the complete and concurrent shut-down of proliferation programs which was only detectably evoked with concentrations of E/F less than or equal to 10 ng/ml. Under these low E/F conditions, mixed glial populations containing only mitotically inactive astrocyte and oligodendroglia lineage cells were generated suggesting that it is not possible to derive actively proliferating immature glia from NPC simply by making small reductions in E/F concentration.

### 3.3. Morphogen directed astroglial differentiation of NPC *in vitro* also requires concurrent cessation of proliferation programs

Since making small reductions to E/F concentrations failed to derive proliferating immature astroglia from cultured NPC, we next probed whether directed differentiation using molecules known to promote astrocyte differentiation *in vitro* could be used to derive astrocytes with maintained proliferation capacity. We first tested the exposure of NPC to either CNTF (100 ng/ml), 1% FBS, or combinations of the two morphogens for 4 days in the absence of E/F (Figure 4A). CNTF and FBS were selected for studying astrocyte differentiation because they are established mediators of astrocyte maturation and contain molecules that act on independent and synergistic signaling pathways^58^. Host astrocytes *in vivo* also upregulate proliferating, immature-like wound repair programs upon exposure to similar cytokine and serum molecules following CNS injury^59,60^. CNTF, FBS, and the combination treatment significantly increased the generation of Gfap-positive astrocytes compared to the SPONT differentiation condition such that FBS containing conditions evoked Gfap expression in 91.8% of cells while treatment with CNTF alone triggered Gfap expression in 72.8% of cells which was slightly, but significantly, less than FBS containing conditions (Figure 4B, C). Gfap-positive cell morphology varied depending on the morphogens used, with 1% FBS treated cells having flat and polygonal shaped phenotypes, while cells exposed to CNTF-only were observed as multicellular clusters with thin, protruding Gfap-positive filaments. The combination of FBS and CNTF treatment resulted is cells with an intermediary phenotype between those observed for the individual morphogens leading to flattened cells that extended many fine Gfap-positive processes (Figure 4B). The increased number of Gfap-positive cells stimulated by directed differentiation coincided with a significant decrease in the proportion of Pdgfra-positive oligodendroglia suggesting preferential astrocyte differentiation under these directed differentiation conditions (Figure 4C). For all directed differentiation conditions, just like for SPONT differentiation, there was a comprehensive loss of Top2a and Ki-67 protein expression such that less than 5% of cells still had detectable levels of these proliferation markers at 4 days and gene expression levels of *Top2a, Mki67* and *Igfbp2* were downregulated by as much as 90-fold (Figure 4D-F, S3C). The reduction in proliferation capacity in directed differentiated cells also lead to a 71-89% decrease in total seeded cell numbers by 2 days of culture that was maintained essentially unchanged by 4 days (Figure S3A). All directed differentiation conditions yielded mitotically inactive, quiescent cells.

**Figure 4.**
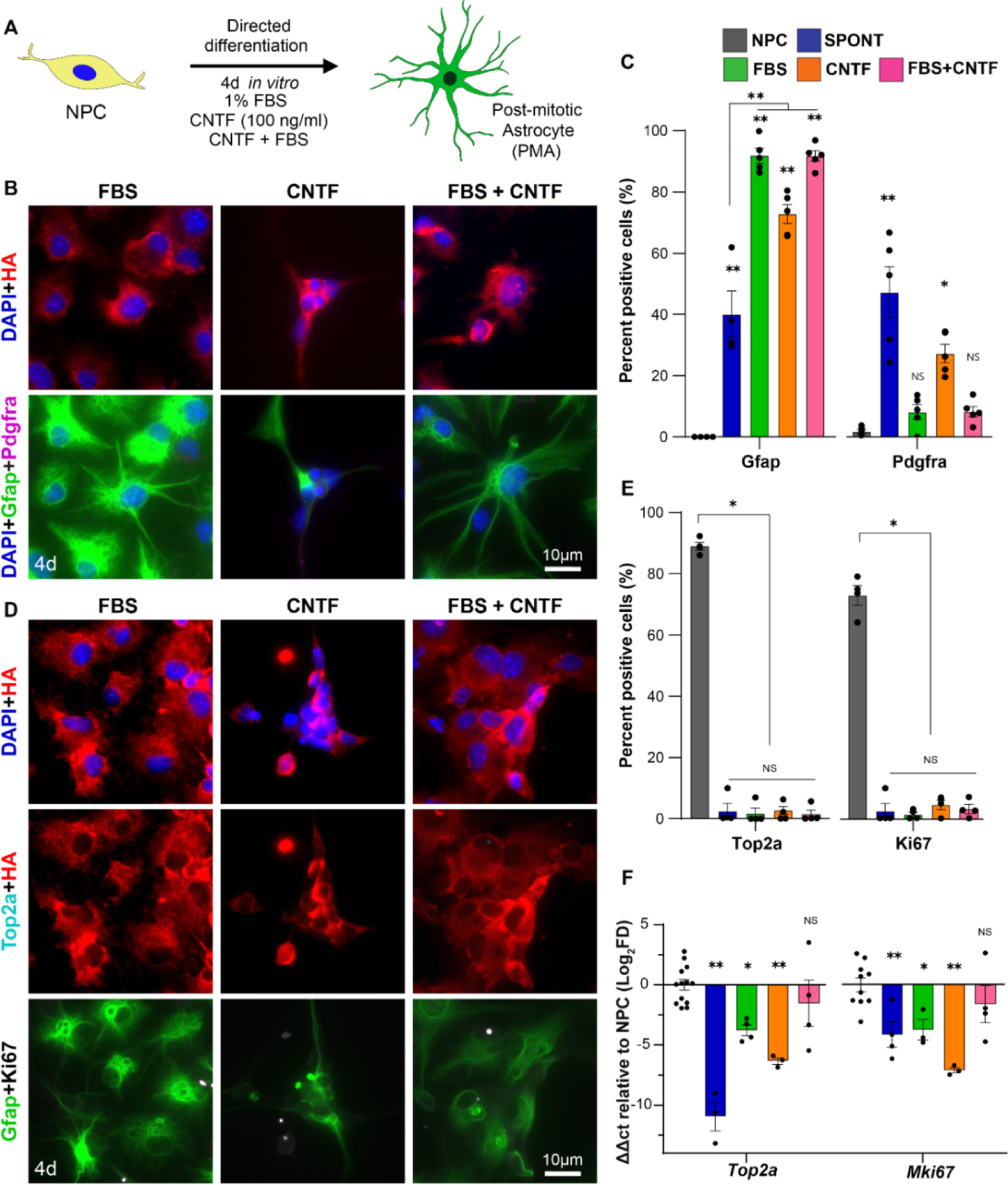
Directed differentiation of NPC yield predominantly quiescent astrocytes. **A.** Schematic of experimental approach for studying directed differentiation using FBS, CNTF, both (CNTF+FBS). **B.** ICC detailed images of directed differentiation of HA-positive NPC into predominately Gfap-positive astrocytes and minimal Pdgfra-positive oligodendroglia. **C.** Quantification of the proportion of astrocytes (Gfap) and oligodendroglia (Pdgfra) derived from directed differentiation with CNTF and FBS morphogens. One-way ANOVA with Tukey test, not significant (NS), * p-value < 0.003, ** p-value < 0.0001. Individual data point showing n=5 for all groups. Sample are compared to NPCs at 100 ng/ml (Grey). **D.** ICC detailed images showing directed differentiation-induced suppression of proliferation markers (Top2a and Ki67). **E.** Quantification of the proportion of differentiated cells expressing proliferation markers (Top2a and Ki67). One-way ANOVA with Tukey test, not significant (NS), * p-value < 0.0001. Individual data point showing n=4 for all groups. Sample are compared to NPCs at 100 ng/ml (Grey). **F.** Directed differentiation effects on *Top2a* and *Mki67* gene expression by qPCR. ΔΔct values normalized to NPC (100 ng/ml E/F) with *Rpl22-HA* used as a housekeeping gene. One-way ANOVA with Tukey test, not significant (NS), * p-value < 0.03, ** p-value < 0.0005. Individual data point showing n=13 for NPC *Top2a*, n=9 for NPC *Mki67*, n=3 for SPONT *Top2a*, FBS *Mki67*, CNTF *Top2a*, CNTF *Mki67* and n=4 for all other groups. All graphs are mean±s.e.m. Sample are compared to NPCs at 100 ng/ml (Grey).

Next, we tested whether concurrent exposure to these directed differentiation morphogens along with maintained high concentrations of E/F (100 ng/ml) could lead to generation of proliferating astroglia as we rationalized that there could be additive effects from both cues (Figure S4). After 4 days of treatment with astrocyte directed differentiation morphogens and E/F mitogens together, we surprisingly detected no Gfap-positive cells using either CNTF or 1%FBS (Figure S4A-B). Interestingly, while CNTF treated cells had the same number of Top2a and Ki67 positive cells as regular NPC cultures, the FBS treated samples, even with the addition of high E/F treatment, had reduced expression of both Top2a and Ki67 suggesting that serum contains potent cytostatic or anti-mitotic cues (Figure S4B). Similar inhibition of proliferation in spite of the presence of E/F mitogens has been noted in neonate astrocytes co-treated with bone morphogenic proteins (BMP)^61^ and FBS notably contains such cytokines^62,63^. Concurrent FBS and E/F treatment dramatically altered cell morphologies compared to NPC while the CNTF plus E/F treatment induced no notable phenotype change (Figure S4A). Although maintaining high concentrations of E/F concurrently with directed differentiation morphogens did not confer expression of the Gfap protein at detectable levels, there was some small but significant increase in Gfap transcripts detected by qPCR in both CNTF and FBS treated samples (Figure S4C).

These data demonstrate that driving glial cell differentiation of NPC *in vitro*, even when employing potent differentiation morphogens, requires cell cycle exit that can only be achieved by reducing culture media concentrations of E/F mitogens. These results provide compelling evidence that for NPC, cell proliferation and differentiation have a strong inverse dependency in a manner consistent with other cell-type progenitors^64^ and mirrors the *in vitro* maturation of primary neonatal astrocytes ^65^.

### 3.4. Hysteretic conditioning of NPC *in vitro* generates proliferating immature astrocytes

Given that single culture medium formulations failed to generate immature astrocytes directly from NPC, we next explored the possibility of deriving immature astrocytes using a sequential approach. This sequential approach was inspired by the concept of hysteresis, which is a phenomenon observed across engineering systems, such as magnetic materials, whereby the current state of the system is dependent not only on the present stimulus but on its recent history^66^. Hysteresis phenomena applied to biology has been used to describe cellular phenotypic memory^67^, and has been exploited in the development of sequential treatments to overcome antibiotic resistance in bacteria^68^. Our strategy for hysteretic conditioning (HC) of NPC explored here involves cyclic, transient exposure of NPC to differentiation directing morphogens to induce astrocyte differentiation (AD) followed by re-exposure to high concentrations of E/F to reinvigorate cell proliferation and stimulate immaturity (SI) programs (Figure 5A). We hypothesized that such sequential treatment would result in hysteresis of key astrocyte characteristics in these cells such that proliferating immature astroglia would be derived, rather than a reversal back to an original NPC state, upon re-exposure to high E/F concentrations (Figure 5A). To test this strategy, we first exposed NPC to the three directed differentiation morphogen formulations tested previously, namely 1% FBS, CNTF, and 1%FBS+CNTF. Since a quiescent astrocyte state is induced by these morphogens by 2 days and changes minimally with 2 additional days of treatment (Figure S5), we chose to expose cells to directed differentiation conditions for the shorter duration to enable an efficient derivation procedure. After 2 days of directed differentiation, cells were re-exposed to high concentration E/F (100 ng/ml) for 3 days and then evaluated by ICC and qPCR. This 2+3 procedure represented one complete cycle of HC. Remarkably, all groups of HC-cells, irrespective of the directed differentiation morphogens used, maintained Gfap expression in 94.5% of cells at the end of 2+3 HC while also showing concurrently high co-expression of Top2a and Ki67 at levels that were not different to that detected in NPC (Figure 5B-E). Gfap-positive HC-cells also robustly co-expressed the immature astrocyte marker, Vimentin (Vim), whereas quiescent astrocytes derived by directed differentiation only did not (Figure 5A-B, S6A). HC conditioning groups selectively enriched for immature astroglia by depleting the proportion of Pdgfra-positive oligodendroglia lineage cells and suppressing the gene expression of *Tnr* compared to SPONT differentiation (Figure 5D, S6). Although nearly every cell expressed high levels of Gfap protein across all HC methods, the relative *Gfap* transcript levels were 18-fold lower compared to the expression in cells after 2 days of directed differentiation (Figure 5B-D, S6B). Since *Gfap* gene expression levels are strongly correlated with maturation state^18^, these data suggest HC-cells are less mature astrocytes than the quiescent astrocytes derived by directed differentiation alone. Expression of proliferation genes *Top2a* and *Mki67* as well as the immaturity gene *Igfbp2* were maintained at NPC-like levels across all the different HC-cells (Figure 5E, S6C-D). Notably, at the end of the AD stage, HC-cells showed complete cell cycle arrest comparable to the quiescent astrocytes, but after the SI stage HC-cell growth rates were no different from those maintained by NPC (Figure 5F).

**Figure 5.**
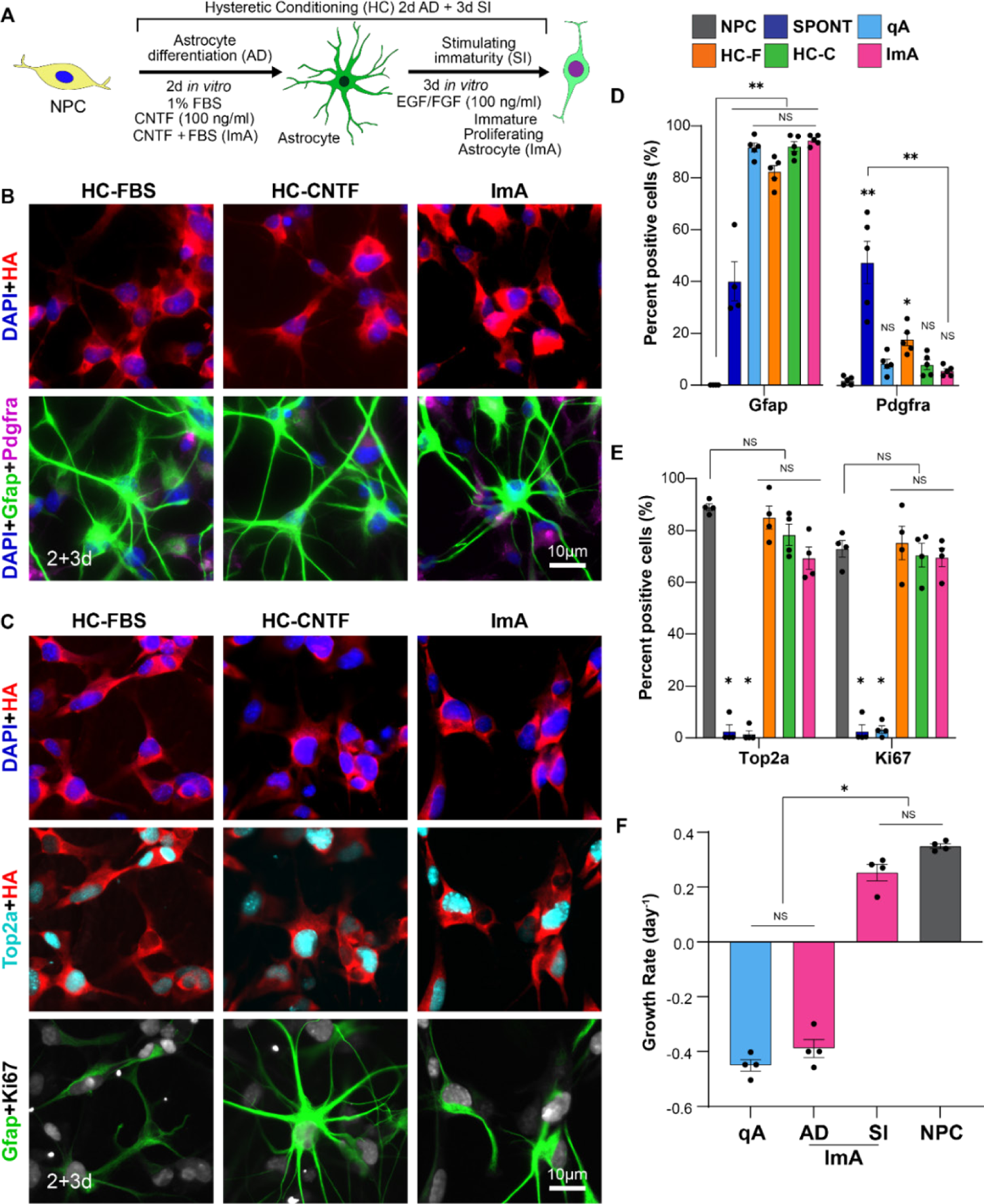
Hysteretic conditioning (HC) of NPC generates astrocytes with high proliferation capacity. **A.** Schematic outlining the hysteretic conditioning approach that involves 2 days of astrocyte differentiation (AD) and 3 days of stimulating immaturity (SI) with 100 ng/ml EGF/FGF (E/F). **B&C.** ICC detailed images after hysteretic conditioning of NPC with FBS (HC-FBS), CNTF (HC-CNTF) or CNT+FBS (ImA) showing the generation Gfap-positive astrocytes (**B.**) that robustly express proliferation markers Top2a and Ki67 (**C**.). **D.** Quantification of the proportion of astrocytes (Gfap) and oligodendrocytes (Pdgfra) derived from HC. One-way ANOVA with Tukey test, not significant (NS), * p-value < 0.05, ** p-value < 0.0001. Individual data point showing n=5 for all groups. Sample are compared to NPCs at 100 ng/ml (Grey). **E.** Quantification of the proportion of cells expressing proliferation markers (Top2a and Ki67) following HC. Two-way ANOVA with Tukey test, not significant (NS), * p-value < 0.0001. Individual data point showing n=4 for all groups. Sample are compared to NPCs at 100 ng/ml (Grey). **F.** Cell number exponential growth rates for qA, ImA at the end of AD and SI phases of the HC, and NPC. One-way ANOVA with Tukey test, not significant (NS), * p-value < 0.0001. Individual data point showing n=4 for all groups. All graphs are mean±s.e.m.

We next tested whether increasing the number of cycles of HC would alter the phenotype of HC-cells. Cells exposed to 2 or 3 cycles of HC showed no significant differences in the expression of *Gfap, S100b, Tnr,* or *Igfbp2, Top2a* and *Ki67* proliferation genes, but there was a small but significant increase in expression of an established astrocyte reactivity marker^18^, *Emp3*,with increasing cycle number as well as a slight reduction in *Gfap* and *Tnr* expression after 3 cycles (Figure S6A, B). Since there were minimal changes in HC-cells with additional treatment cycles we opted to pursue one HC cycle only for immature astrocyte derivation for the subsequent evaluations to enable an efficient derivation procedure. We then tested how prolonged E/F exposure altered the astrocyte hysteresis by keeping HC-cells in high concentration (100 ng/ml) E/F media for 21 days. Compared to the 3 days of E/F exposure, we detected significant decrease in *Gfap* and significant increase in *Top2a* and *Mki67* gene expression suggestion a progressive loss of astroglial phenotype and reversion towards an NPC-like state with more prolonged culture with E/F mitogens (Figure S6C, D). In considering all the above data collectively, we chose to use a 2+3 day hysteretic conditioning paradigm using the combined CNTF+FBS treatment for directed differentiation to derive the HC-cells used for subsequent experiments in this study. These selected HC-cells are referred to henceforth as immature astrocytes (ImA).

To evaluate whether ImA had altered cell autonomous differentiation capacity, we next evaluated the phenotype of ImA following 2 and 4 days of E/F withdrawal. Under these SPONT differentiation conditions, there was no difference in the expression of *Gfap*, *Tnr* or *Emp3* trancripts for progeny derived from either ImA or NPC (Figure S8A). However, differentiation of ImA generated a significant increase in the proportion of Gfap-positive cells and a concurrent decrease in Pdgfra-positive cells by ICC, compared to SPONT differentiation of NPC (Figure S8C). Gfap-positive progeny derived from ImA expressed Gfap at a higher intensity per cell compared to Gfap-positive cells derived from NPC (Figure 8C). There was no detectable expression of Top2a or Ki67 in ImA following withdrawal of E/F by ICC and gene expression of *Mki67, Top2a* and *Igfbp2* were downregulated to similar levels as NPC following differentiation suggesting that the capacity to enter a state of quiescence was not altered by the HC procedure.

These data show that hysteretic conditioning of NPC by sequential directed astrocyte differentiation and immaturity stimulation derives a population of proliferating ImA after one HC cycle. ImA are cell autonomously primed to spontaneously differentiate along an astroglial lineage upon E/F removal which improves on the heterogenous mixed glial cell differentiation observed for spontaneous differentiation of NPC. Overall, these data demonstrate that HC is a viable strategy for deriving enriched populations of proliferating ImA for grafting and are examined for such a purpose in comparison with NPC in more detail below.

### 3.5. Immature astroglia are more resistant to myofibroblast-like transitions than NPC *in vitro* but maintain wound healing migration capacity

High concentration serum environments, such as those present in CNS injury lesions, non-cell autonomously modify NPC by driving myofibroblast-like phenotypes which may reduce glia repair capacity^18^. To test whether ImA are differentially altered by such high concentration serum conditions compared to NPC, we evaluated cells after 4 days exposure to media enriched with 5% FBS. Under these conditions there is a notable epithelial-mesenchymal transition (EMT)-like effect on NPC that provokes a defined myofibroblast-like phenotype including increased expression of canonical EMT gene *Acta2*, synthesis and assembly of contractile actin stress fibers, and increases in cytoplasm and nucleus size^18^ (Figure 6A, S9). To quantify the overall extent of these EMT-like effects, a multifactorial EMT score was derived from a principal component analysis (PCA) that incorporated data from numerous relevant evaluations of cell phenotype made using ICC and qPCR readouts (Figure 6B, S9). Exposure to serum enriched media induced a significant increase in EMT score for both NPC and ImA compared to their pre-exposure baselines. However, compared to NPC, ImA had a reduced EMT score reflecting a small but significant attenuation of the myofibroblast-like transition effect under serum enriched environments (Figure 6B-D, S9). Compared to serum exposed NPC, the ImA had: (i) reduced upregulation of *Acta2*; (ii) increased overall expression of Gfap; (iii) reduced expression of F-Actin stress fibers that are characteristic of a myofibroblast-like phenotype; and (iv) a significantly smaller increase in cytoplasm and nucleus size (Figure S9).

**Figure 6.**
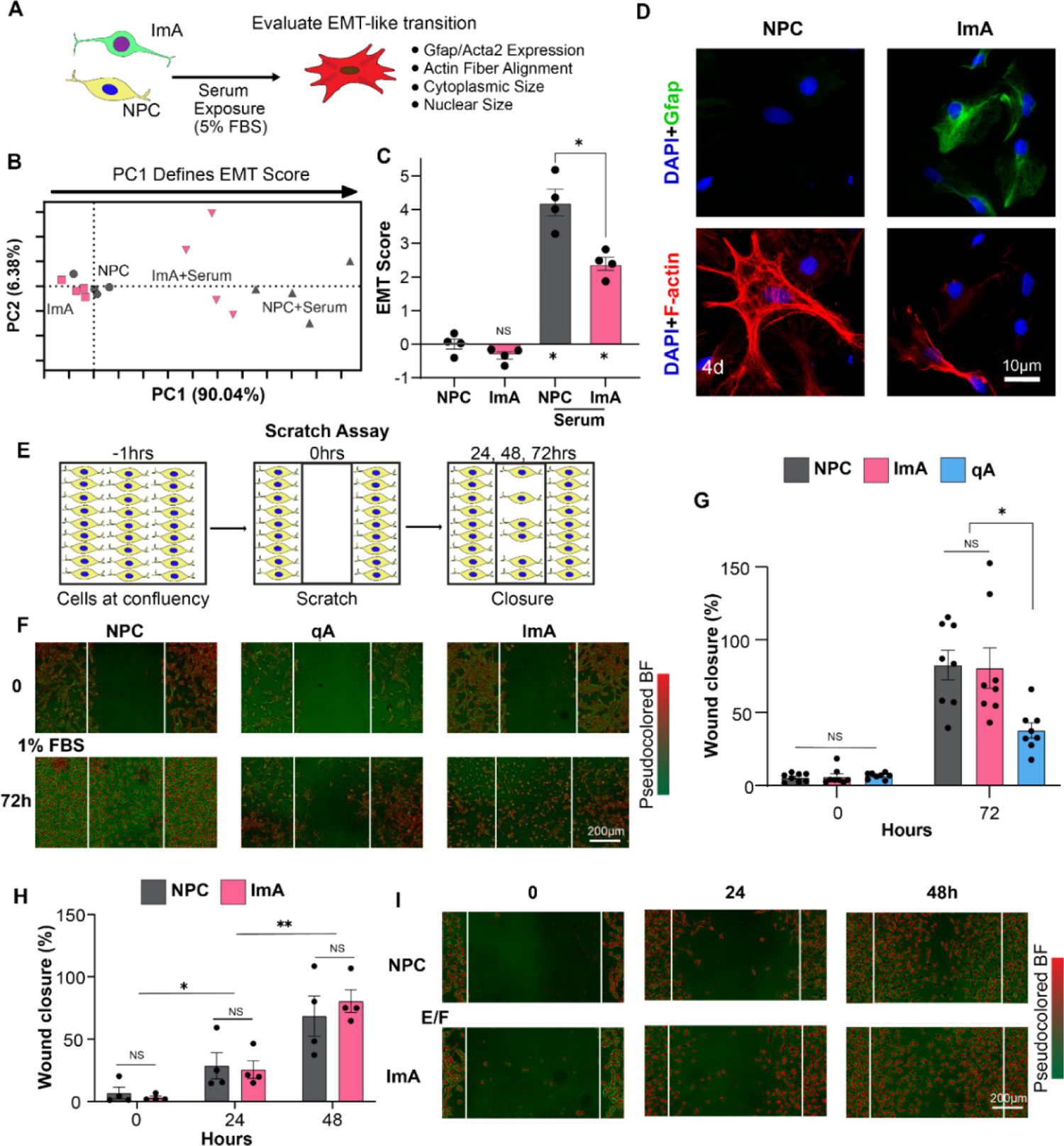
ImA are more resistant to myofibroblast-like (EMT-like) transitions than NPC *in vitro*. **A.** Schematic of experimental approach assessing ImA and NPC following 5% serum (FBS) exposure. **B.** PCA incorporating multiple ICC and qPCR-based evaluations of EMT-like effects showing differences in how ImA and NPC respond to serum exposure. **C.** Comparison of EMT score derived from PC1 of PCA for NPC and ImA before and after serum exposure. Two-way ANOVA with Tukey test, * p-value < 0.0001. Individual data point showing n=4 for all groups. **D** ICC detailed images of NPC and ImA before and after serum exposure, stained with Phalloidin (F-Actin), which shows the extent of the myofibroblast-like phenotype, and Gfap. **E.** Schematic of experimental approach for wound healing scratch assay. **F.** Pseudocolored brightfield images of NPC, qA, and ImA at scratch region at 0 and 72 hours in 1% FBS containing media. **G.** Quantification of scratch wound closure for NPC, qA and ImA at time of scratch and 72 hours after scratch. Two-way ANOVA with Tukey test, not significant (NS), * p-value < 0.0005. Individual data point showing n=8 for all groups. **H.** Quantification of scratch wound closure for NPC and ImA with addition of EGF/FGF mitogens to the media. Two-way ANOVA with Tukey test, not significant (ns), * p-value < 0.04, ** p-value < 0.002. Individual data point showing n=4 for all groups. **I.** Pseudocolored brightfield images of NPC and ImA responses scratch region at 0, 24 and 48 hours after scratch with E/F treatment. All graphs are mean±s.e.m.

Although engagement of EMT-like programs may perturb glia repair competency^18^, it can be an important and necessary response in cells adjacent to tissue injury to enable otherwise non-motile cells to migrate across a wound site^69^. Since this competency is likely to be important for wound repair functions of grafts^69^, we next tested whether an attenuation of the myofibroblast-like responses in ImA caused a concurrent reduction in effective cell migration capacity. To evaluate cell migration, we used a standardized 2D monolayer scratch assay and evaluated migration of ImA, NPC, and qA over a 72-hour period after scratch (Figure 6E).

Although cell monolayers were first prepared in their own appropriate media for that group, after scratch, all groups were maintained in 1% FBS without E/F growth factors to simulate a CNS injury lesion environment and minimize the contribution of cell proliferation on the re-population of cells in the scratched zone. While qA showed a relatively poor capacity to fill in the scratched zone resulting in 38% filling of the scratched area by migrating cells over 72 hours, there was a notably higher cell migration capacity in ImA and NPC with both cell types recapitulating more than 80% of the pre-scratch cell density by 72 hours (Figure 6F, G). When provided media containing E/F, the NPC and ImA were also both able to recapitulate a similar cellular density within the injury zone as early as 48 hours following scratch, reinforcing the importance of growth factor mitogen signaling for also enabling efficient cell migration in wound repair (Figure 6H, I).

These data show that compared to NPC, ImA derived by hysteretic conditioning are less susceptible to non-cell autonomous cues from serum enriched environments *in vitro* that otherwise direct cells away from astrocyte differentiation and towards myofibroblast-like phenotypes. Importantly for wound repair functions, the migration capacity of ImA following an *in vitro* wound was comparable to NPC and is unaffected by its capacity to attenuate EMT-like transformation.

### 3.6. Immature astroglia generated by hysteretic conditioning direct comparable glia repair as NPC when grafted into ischemic strokes

Since ImA showed attenuated EMT-like phenotypes in high serum environments *in vitro* compared to NPC, we hypothesized that ImA may have an improved capacity to maintain wound repair astrocyte phenotypes upon grafting into severe CNS injury lesions. To evaluate ImA grafts, we began by comparing outcomes following grafting into healthy striatum (Figure S10) and striatal ischemic stroke lesions (Figure 7) and benchmarking ImA performance against NPC grafts which we have evaluated extensively in our previous work^18^. Grafts of quiescent astrocytes (qA) derived by 4-day directed differentiation with FBS+CNTF were also tested to assess the effect of proliferating versus nonproliferating astrocyte grafts on glia repair outcomes. Across all studies, grafts were prepared using supplemented media but without the aid of any other transplantation carrier. Tissue sections were evaluated by IHC to identify the phenotypes of HA-positive graft progeny at 3-and 12-days post transplantation (Figure 7A-B). When grafted into healthy striatum, ImA survived at greater numbers than qA resulting in recruitment of greater numbers of host astrocyte and oligodendroglia to the graft site (Figure S10). The ImA grafts remained localized within a radial distance of 200 µm from the site of injection, with essentially all ImA graft-derived progeny expressing Gfap with no Pdgfra-positive oligodendroglia lineage cells being detected. By contrast and as noted before^18^, NPC grafts in healthy neural tissue generate numerous Pdgfra-positive oligodendroglia but such cells tend to only be detected in neural tissue regions at least 300 µm away from the injection epicenter and we did not detect any ImA progeny at such distances.

**Figure 7.**
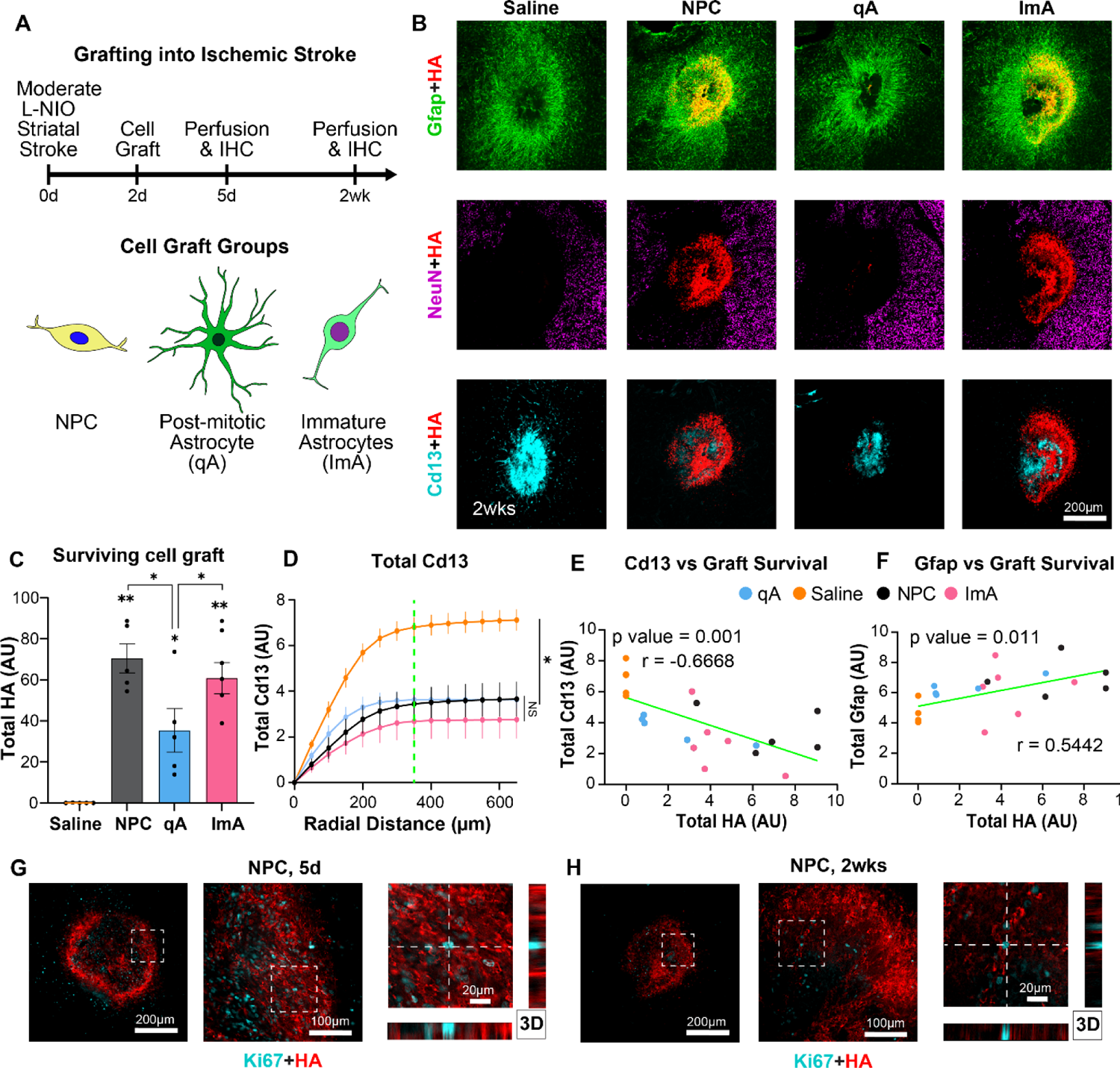
Comparing cell grafts in ischemic strokes. **A.** Schematic summarizing experimental details of NPC, ImA and qA grafts into L-NIO induced striatal ischemic stroke. **B.** Survey IHC images at 2 weeks post stroke showing survival of HA-positive grafts, astrocyte responses (Gfap), non-neural lesion cores (Cd13) and regions of preserved neural tissue (NeuN). **C** Quantification of total HA-positive grafted cell number at stroke lesions. One-way ANOVA with Tukey test, * p-value < 0.05, ** p-value < 0.0001. Individual data point showing n=5 for saline, NPC and qA and n=6 for ImA. Samples are compared to Saline at 100 ng/ml (Orange). **D.** Quantification of Total Cd13 radially from center of the lesion at 50 μm intervals calculated as area under the curve (AUC) of the radial Cd13 intensity plot. Green line indicates border of non-neural lesion. One-way ANOVA with Tukey test, not significant (NS), * p-value < 0.0002. n=5 for saline, NPC and qA and n=6 for ImA. **E.** Correlation analysis of total Cd13 with non-neural lesions compared with total HA-positive grafted cells showing an inverse correlation. Individual data point for each animal showing n=5 for saline, NPC and qA and n=6 for ImA. **F.** Correlation analysis of total Gfap within lesions and total HA-positive grafted cells showing positive correlation. Individual data point for each animal showing n=5 for saline, NPC and qA and n=6 for ImA. **G&H.** High magnification 3D IHC images showing no detectable Ki67-positive cells in HA-positive graft progeny at either 5 days (**G.**) or 2 weeks (**H.**). All graphs are mean±s.e.m.

Grafting of NPC, ImA, and qA into striatal ischemic strokes at two days post injury resulted in increased numbers of detectable HA-positive cells for each graft type compared to the same graft made into healthy striatum (Figure 7A-C, S11). Despite the detectable increase in graft-derived HA-positive cells at ischemic stroke lesions, we could not identify any Ki67-positive proliferating graft progeny at either 3 or 12 days after transplantation (Figure 7G, H, S11) despite identifying some Ki67/Sox9/Gfap-positive host astrocytes adjacent to HA-positive grafts at the earlier time point and numerous Ki67/Cd13-positive inflammatory cells in lesion cores at the later timepoint (Figure 7G, H, S11). Notably, as early as 3 days, progeny of NPC grafts were essentially all Sox9- and Gfap-positive suggesting that a rapid astrocyte maturation coincided with the loss of proliferation capacity in grafts which is consistent with the inverse correlation between these two functions observed from the earlier *in vitro* studies. ImA and NPC had equivalent numbers of detected HA-positive cells at ischemic stroke lesions, while both grafts had greater cell survival compared to qA (Figure 7C, S12). All grafts directed significant alterations to the ischemic stroke lesion with HA-positive cells dispersing throughout the lesion core resulting in a significant reduction in the total number of Cd13-positive cells and fibrotic lesion size compared to saline treated strokes (Figure 7D, S12B, F). Across all grafts, the total number of Cd13-positive lesion core cells was significantly inversely correlated (r=-0.67) with grafted cell survival supporting the notion that the grafts contribute directly to the altered wound repair detected (Figure 7D, E, S12). NPC and ImA grafts also contributed to increasing the number of Gfap-positive cells within lesions to alter the location of the astroglial lesion border and increase the density of Cd31-positive angiogenic vessels in lesion cores compared to saline controls (Figure 7F, S13). None of the cell graft groups altered the overall volume of neural tissue damaged by the ischemic stroke and we detected no graft derived neurons in lesions (Figure S12), which was consistent with previous observations^18^

These data show that cell grafts, irrespective of their pre-graft transcriptional state, survive better within an injury environment created by L-NIO ischemic injuries compared to uninjured healthy tissue and suggest that there is enhanced trophic or metabolic support for cell survival within these lesion environments compared to healthy tissue. Furthermore, the extent of glia-based repair and reduction in fibrotic scarring observed in ischemic lesion cores was directly proportional to overall graft cell survival across the three types of grafts. Grafts that were actively proliferating at the time of transplantation, namely NPC and ImA, had the best cell survival and the most effective glia repair outcomes. Notably, despite essentially all NPC and ImA graft cells actively proliferating at the time of transplantation, mitotic activity was suppressed in grafts as early as 3 days post transplantation. The comparatively similar glia repair phenotypes at lesions for both ImA and NPC grafts despite large differences in their pre-graft transcriptional states provides further evidence to the dominant impact of non-cell autonomous regulation of graft cell fate in CNS injury lesions.

### 3.7. NPC and ImA grafts converge to similar phenotypes in hemorrhagic strokes with reduced glia repair capacity compared to ischemic injuries

Given comparable glia repair outcomes between NPC and ImA grafts in ischemic stroke lesions, we decided to further compare the two graft types in collagenase-induced hemorrhagic stroke lesions to test whether ImA performed better in these more severe lesions where there is a greater extent of bleeding and less angiogenesis in lesion cores. As before, we grafted NPC and ImA into hemorrhagic lesions at 2 days after injury and compared outcomes with saline treated controls (Figure 8A). At 2 weeks after injury, we detected low overall cell survival and no notable differences between the NPC and ImA grafts (Figure 8B). For both grafts, HA-positive cells progeny were detected mostly at the host astroglial border margins of hemorrhagic lesions with only a few cells identified near the center of lesions. Compared to the grafts made in ischemic strokes, there were nearly 84% fewer HA-positive cells that survived in hemorrhagic strokes for both NPC and ImA (Figure 8C). The multi-cellular composition of hemorrhagic lesions (Figure 8B, S1), the phenotype of astrocyte borders (Figure 8, S1, S14, S15), and the total numbers of recruited Cd13-positive immune cells was essentially unaltered by either the NPC or ImA grafts compared to saline-treated controls (Figure 8B, D). In lesion volumes where grafted cells did survive at moderate numbers, local Cd13-positive cell density was noticeably, although minimally, decreased (Figure 8B). For both NPC and ImA, there was a small but significant reduction in the proportion of the Gfap-positive grafted cells identified in hemorrhagic lesions when compared to the prominent astroglia phenotypes adopted by grafts in ischemic lesions.

**Figure 8.**
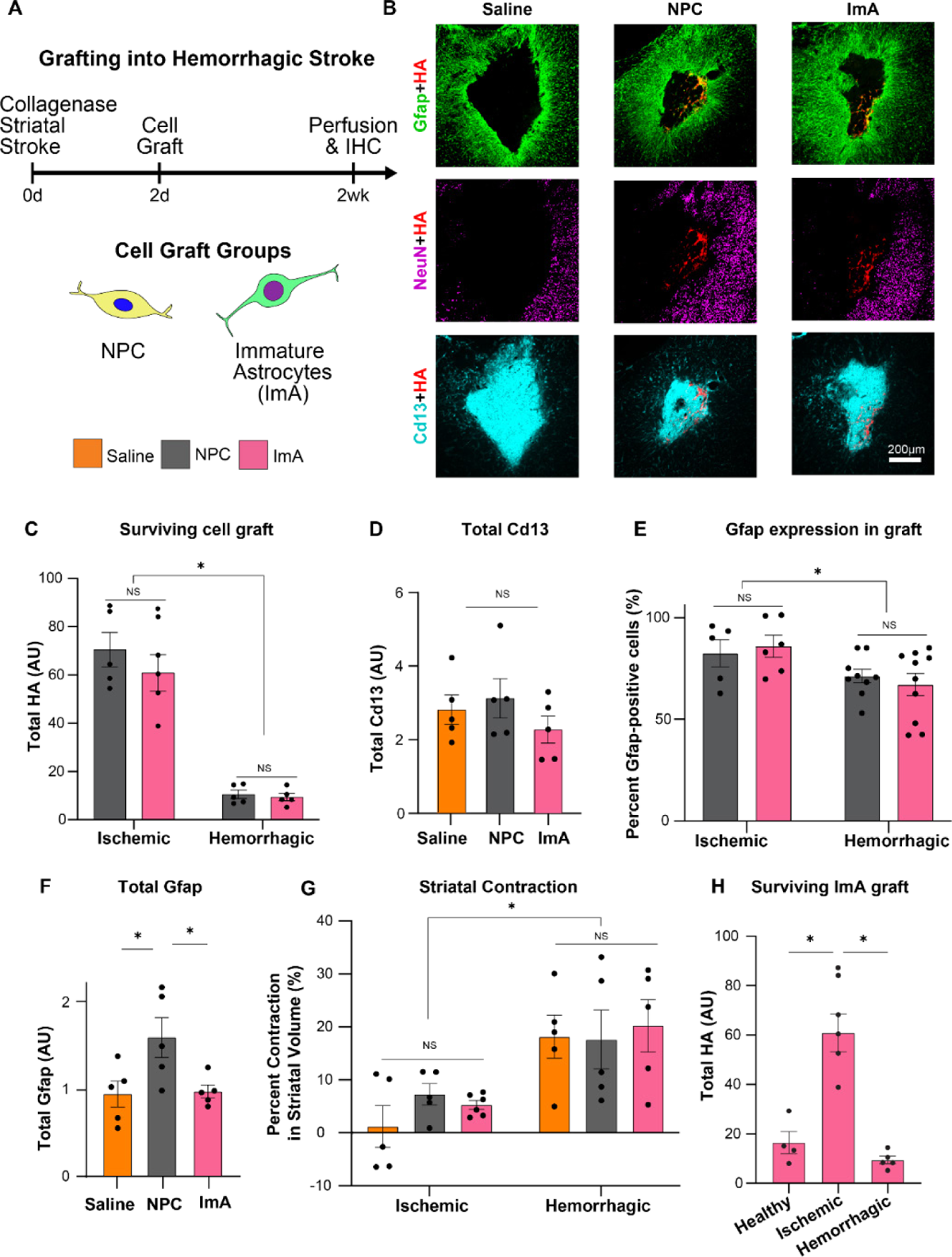
Comparing cell grafts in hemorrhagic stroke. **A.** Schematic summarizing experimental details of NPC, ImA and qA grafts in collagenase induced striatal hemorrhagic stroke. **B.** Survey IHC images at 2 weeks post stroke showing survival of HA-positive grafts, astrocyte responses (Gfap), non-neural lesion cores (Cd13) and regions of preserved neural tissue (NeuN). **C.** Quantification of total HA-positive grafted cell number at stroke lesions. Two-way ANOVA with Tukey test; * p-value < 0.001. Individual data point showing n=6 for ischemic ImA and n=5 mice for all other groups. **D.** Quantification of total Cd13 at hemorrhagic stroke lesions with and without cell grafts. One-way ANOVA with Tukey test, not significant (NS). Individual data points showing n=5 mice for all groups. **E.** Quantification of the proportion of Gfap-positive graft progeny for NPC and ImA in ischemic and hemorrhagic strokes. Two-way ANOVA with Tukey test, not significant (NS), * p-value < 0.01. Individual data points showing individual analyzed images with n=5 for NPC and n=6 for ImA in ischemic strokes, n=9 for NPC and n=10 for ImA in hemorrhagic strokes. **F.** Quantification of total Gfap at lesions. One-way ANOVA with Tukey test, * p-value < 0.05. Individual data points showing n=5 mice for all groups. **G.** Quantification of the contraction in striatal volume normalized to contralateral uninjured striatum for ischemic and hemorrhagic strokes treated with saline, NPC and ImA grafts. Two-way ANOVA with Tukey test, not significant (NS), * p-value < 0.0001; Individual data points showing n=5 mice for all groups. **H.** Quantification of total HA-positive cell number for ImA grafts made into healthy striatum as well as ischemic and hemorrhagic stroke injuries. One-way ANOVA with Tukey test; * p-value < 0.0003. Individual data points showing n=4 mice for healthy, n=6 for ischemic, and n=5 for hemorrhagic. All graphs are mean±s.e.m.

Despite this overall reduction in graft derived astroglia, NPC showed a small but significant increase in total Gfap at hemorrhagic lesions compared to ImA, and ImA had total Gfap levels that were surprisingly no different from saline controls (Figure 8F). In hemorrhagic lesions there was a significant contraction and loss of striatal tissue volume at 2 weeks after injury that was not observed in the ischemic injuries, with the extent of tissue contraction in hemorrhagic strokes being unaltered by either type of cell graft compared to the saline-treated controls (Figure 8G). NPCs and ImA were positive for canonical border forming astrocyte markers such as Id3, S100a6, and Pu.1 consistent with their adoption of a wound repair astroglial phenotype (Figure S14). Neither cell graft induced any altered Cd31-positive angiogenesis at hemorrhagic lesion cores compared to saline treated controls (Figure S15).

These data show that NPC and ImA grafts are equivalently altered by lesion environmental cues present in hemorrhagic- and ischemic-stroke lesions and that the two lesion types dramatically and differentially impact overall graft survival and function. Limited grafted cell survival in hemorrhagic lesions ultimately results in ineffective glia repair irrespective of whether cell grafts, such as ImA, are primed with such functions prior to transplantation. It was notable that graft survival was greatest for all grafts in L-NIO induced ischemic lesions and not healthy neural tissue (Figure 8H), suggesting that certain injury environments, or perhaps the host glial response to injury, provides necessary trophic support to grafts that is otherwise absent from either normal healthy tissue or hemorrhagic lesions. Identifying these trophic factors may hold the key to augmenting graft outcomes in CNS injuries more broadly.

## 4. Discussion

In mammalian neonates, proliferating immature astrocytes direct scar-free parenchymal repair after injury that aids neural circuit regeneration, but this competency is lost with adult development^9,25–29,70–74^. There is the potential to use cell grafting strategies during the acute period of CNS injury to mimic neonate glia-based repair mechanisms to improve adult neural regeneration. However, ensuring robust cell graft survival and appropriately guiding graft cell fates such that they are capable of effective glia repair functions in severe injuries remain outstanding challenges. Here, we validated a new hysteretic conditioning strategy for generating ImA that we anticipated could improve graft-derived glia repair outcomes in severe adult CNS injuries compared to NPC grafts. However, while ImA were more resilient to serum enriched environments *in vitro* when compared to NPC, grafting outcomes for ImA in two types of CNS injury lesions were no different to NPC, demonstrating that lesion environments, rather than transcriptional starting states, predominately determine graft survival, cell-fate, and glia repair competency. The findings of this study have broad implications across several areas include for: (i) our understanding of the relationship between astrocyte maturation state and proliferation capacity in development and injury; (ii) the use of conditioning strategies in cell transplantation; and (iii) our understanding of how CNS injury environments non-cell autonomously direct graft outcomes.

Here we demonstrated both *in vitro* and *in vivo* that astrocyte differentiation from NPC requires the concurrent abatement of cell proliferation. These findings are consistent with the dynamic and tightly controlled suppression of proliferative capacity that accompanies astrocyte maturation during the first week of mammalian postnatal development *in vivo* ^75,76^ and the maturation of primary neonatal astrocytes *in vitro*^65^. By applying temporally controlled hysteretic conditioning to NPC to generate ImA, we have demonstrated here that it is possible to reinvigorate proliferative programs in quiescent astrocytes derived from NPC using established NPC mitogens without disrupting key astrocyte characteristics such as cell morphology and Gfap expression. Reversibility of quiescence has been demonstrated in neonate astrocytes after BMP-induced differentiation *in vitro*^61^ and in astrocyte-like B stem cells found in the forebrain ventricular-subventricular zone (V-SVZ) niche which readily reacquire proliferative capacity when appropriately stimulated during periods of neurogenesis^77,78^. It will be insightful to compare the transcriptional profiles of HC-derived ImA with these other Gfap-expressing progenitors in future work. Given ImA derived here were cell autonomously biased towards astrocyte differentiation with minimal contamination from oligodendroglia-lineage cells, hysteretic conditioning may also prove to be a particularly efficient culturing method for generating large quantities of purified astrocytes for mechanistic *in vitro* studies.

It is also notable that the HC-strategy employed here mimics the CNS injury induced proliferative responses of mammalian adult astrocytes *in vivo* and identifying how transcriptionally similar the HC-derived ImA are to wound repair astrocytes during the active proliferation state will be another important area of future study. Adult astrocytes re-enter the cell cycle transiently as part of adaptive changes that are essential for forming neuroprotective astrocyte borders around non-neural lesion cores^1,7,60,79,80^. Proliferation of adult astrocytes responding to CNS injury is temporally and spatially restricted because access to the appropriate concentrations of growth factor mitogens is tightly regulated and gradients of blood derived proteins from lesion cores act as forceful cytostatic factors^81–83^. Our findings reinforce the significant effects that these lesion-derived factors have on the proliferation state of astrocytes and neural progenitors. Both ImA and NPC grafts showed rapid suppression of proliferation marker expression after grafting *in vivo* into stroke injuries which occurred well before host border forming astrocytes responding to injury ceased their period of self-renewal. These results suggest that grafts made into lesion core environments may be receiving different concentrations of growth and cytostatic factors compared to the proliferating host astrocytes that persist in the preserved neural tissue adjacent to the injury zone or that these factors regulate grafted cell functions in divergent ways. The results could also suggest that metabolic support to sustain graft proliferation in lesion environments is inadequate^84^. Developing ways to prolong the delivery of growth factors and inhibition of cytostatic factors at lesions as well as providing sustained metabolic support to grafts will likely be crucial for augmenting proliferative capacity in these cells necessary for effective glia repair.

Conditioning therapy is used routinely prior to hematopoietic cell transplantation to minimize barriers to successful engraftment, but this type of conditioning involves treating the host with a regime of chemotherapy, monoclonal antibody therapy, and/or radiation and not altering the state of the graft^85^. Our study extends the principles of conditioning to the cell graft itself as a way to potentially pre-program graft functions before transplantation. The HC-strategy we developed here represents a new way to approach manufacturing cell therapies that may be broadly applicable for deriving committed progenitors from multi- or pluri-potent cells that are more amenable to scaled-up production. While our focus here was to use a cell conditioning strategy to generate a specific cell phenotype for grafting (i.e. proliferating immature astrocytes), it is intriguing to consider the development of other pre-graft conditioning strategies as part of future work to equip cells with a capacity to thrive in hostile tissue injury environments, such as by exposing cells transiently to sub-threshold hypoxic, inflammatory, or nutrient deficient conditions prior to grafting^86^.

The new findings in this study also advance our understanding of how CNS injury environments direct graft cell fate and build on the foundations of our prior work in some important ways^18^. Here we observed that different types of injuries made in the same anatomical location in the brain can differentially alter graft outcome. Remarkably, our results suggest that L-NIO induced ischemic stroke injuries create lesion environments enriched with some essential trophic support since we detected a consistent improvement in cell graft survival compared to healthy tissue irrespective of the pre-graft transcriptional state of the cells that were transplanted. While it has been known since the early peripheral nerve graft studies of Tello that integration of grafted cells with the host can be improved by making minor neural tissue disruptions at the site of transplantation ^87^, our study shows that cell graft survival may also be improved by transplanting cells into certain types of lesion environments during the acute stages of injury.

The cell grafting field has long resisted grafting into acute lesions because the wound environment has been considered hostile towards grafts^14,88,89^. Our results show that hemorrhagic lesions are indeed particularly hostile to acute grafting whereas ischemic injuries are not. We documented notable spatial and temporal differences in both angiogenesis and host glia responses between ischemic and hemorrhagic lesions that could contribute to these differential grafting outcomes since vascularization is essential for graft survival^90^ and glia responses to injury have contributed to improving the survival of other types of neuronal grafts^91,92^. Future work will need to dissect how key lesion differences between the two injury modes impact grafting outcomes as well as systematically identifying the optimal post-injury grafting timepoint for each type of stroke injury to further clarify the key determinants of graft survival and glia repair functions. Biomaterial-based carriers are also emerging as a promising strategy for enhancing graft cell survival during transplantation and for controlling differentiation of grafted progenitors *in situ*^93^. Based on the differential grafting outcomes observed here it would seem pertinent to explore the incorporation of biomaterials that are capable of augmenting angiogenesis^94^ and/or modulating host glia responses^95^ as carriers for NPC or ImA as a way to improve grafting outcomes in severe hemorrhagic injuries.

In conclusion, we developed an innovative method to generate immature astrocytes from NPC using a sequential hysteretic conditioning strategy. ImA showed a greater resistance to EMT-like transitions compared to NPC and effective scratch wound closure *in vitro*. Both NPC and ImA promoted effective glia repair of ischemic strokes but showed reduced survival and ineffective lesion remodeling in hemorrhagic strokes. The results from this study lay the foundation for further investigation into the mechanisms associated with lesion dependent non-cell autonomous regulation of cell graft fate, and prompt renewed efforts to bioengineer approaches to support graft metabolic and trophic demands and mitigate non-cell autonomous regulators present in lesion environments to improve graft survival and functions in severe CNS injuries.

## Acknowledgements

This work was supported by internal Boston University start-up funds (T.M.O), and the Craig H. Neilsen Foundation (732495 to T.M.O.). H.O.A was funded by a Boston University Biomedical Engineering Distinguished Fellowship and Faculty for the Future from Schlumberger Foundation. This research was supported by the Biomedical Engineering Core Facilities at Boston University. A special thanks to Xin Brown and the Biointerface Technologies (BIT) core, as well as the Micro and Nano Imaging (MNI) core at BU.

## Conflict of Interest

The authors declare no conflict of interest.

## Data availability statement

All data generated for this study are included in the main and supplementary figures. For all quantitative figures, the results of statistical tests are provided with the paper. Files of source data of individual values and other data that support the findings of this study are available on reasonable request from the corresponding author.

## Notes

### Competing Interest Statement

The authors have declared no competing interest.

