## Supplementary Information for "Ischemic and hemorrhagic stroke lesion environments differentially alter the glia repair potential of neural progenitor cell and immature astrocyte grafts"

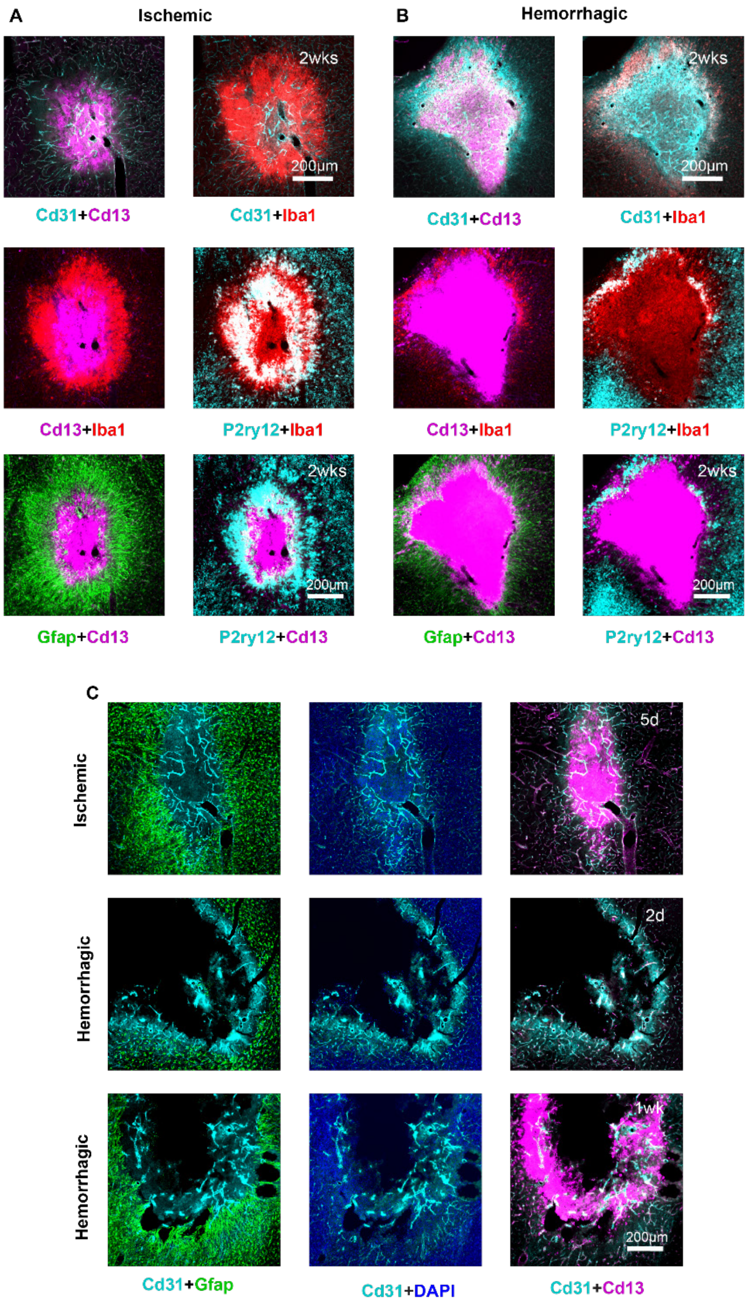


**Supplementary Figure 1**. Ischemic and hemorrhagic stroke have different astroglia, microglia and angiogenic response. **A&B** IHC of ischemic (B) and hemorrhagic (C) stroke at 2 weeks showing platelet endothelial cell adhesion molecule (Cd31), myeloid lineage cells (Cd13), microglia (Iba1, P2ry12), and astrocytes (Gfap). Showing difference in vasculature, microglia response and astroglia border density. **C** IHC of ischemic stroke at 5 days and hemorrhagic stroke at 2 days and 1 week showing astrocytes (Gfap), nucleus (DAPI), platelet endothelial cell adhesion molecule (Cd31) and myeloid lineage cells (Cd13) showing that angiogenesis occurs earlier in ischemic strokes.

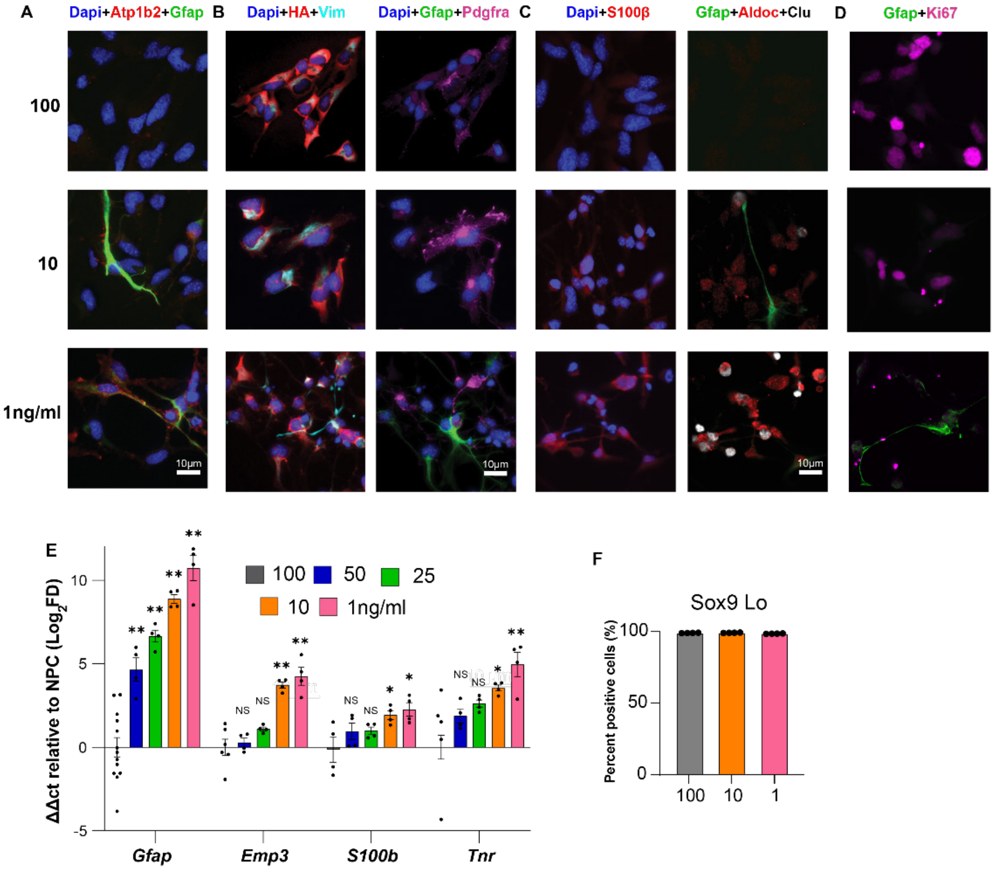


**Supplementary Figure 2.** Reduced concentrations of EGF/FGF(E/F) yield astroglia cells. **A** ICC staining of E/F concentration based spontaneous differentiation of NPC for astrocyte (Gfap, Atp1b2) and nucleus (DAPI). **B** ICC staining of E/F concentration-based spontaneously differentiated NPC for Ribo Tag (HA), nucleus (DAPI), astrocytes (Gfap andVimentin), and oligodendrocytes (Pdgfra). **C** ICC staining of E/F concentration-based spontaneously differentiated NPC for nucleus (DAPI), astrocytes (Gfap, s100b) and reactive astrocytes (Aldoc and Clusterin). **D** ICC staining of E/F concentration-based spontaneously differentiated NPC for nucleus (DAPI), astrocytes (Gfap) and proliferation marker (Ki67). **E** qPCR of astrocyte (*Gfap, S100b*), reactive astrocytes (*Emp3*) and oligodendrocyte (*Tnr*) genes. ΔΔct values normalized to NPC (100 ng/ml E/F) with *Rpl22-HA* used as a housekeeping gene. One-way ANOVA with Tukey test, not significant (NS), * p-value < 0.05, ** p-value < 0.0006. Individual data point showing n=13 for NPC and n=4 for all other groups.. **F** Quantification of percentage positive Sox9Lo Cells. All graphs are mean±s.e.m.

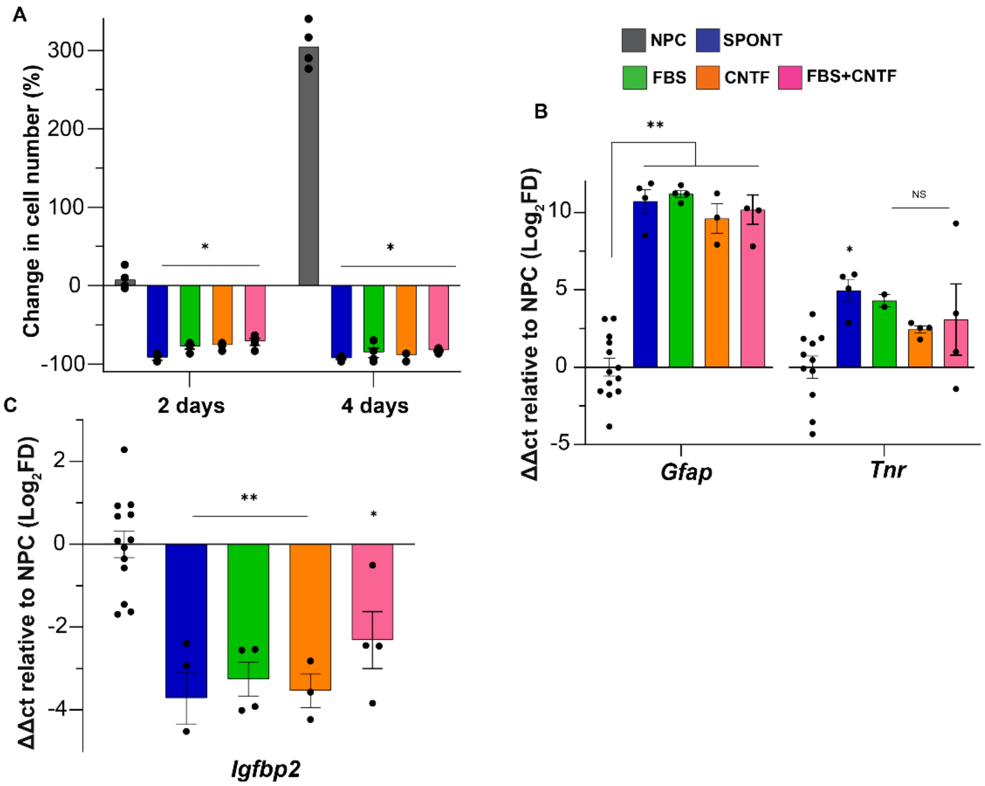


**Supplementary Figure 3.** Directed differentiation of NPCs loss proliferative and immaturity capacity but increase astrocyte expression. **A** Percentage change in cell number of NPC, SPONT (spontaneously differentiated NPC with 1 ng/ml E/F), 1% FBS 100 ng/ml CNTF and 1%FBS and 100 ng/ml CNTF (FBS+CNTF) at 2 and 4 days. Graph shows mean± s.e.m, two-way ANOVA with Tukey test, *** p-value < 0.0001. Individual data point showing n=4 for all groups. **B** qPCR of astrocyte (*Gfap*) and Oligodendrocyte (*Tnr*) genes. ΔΔct values normalized to NPC (100 ng/ml E/F) with *Rpl22-HA* used as a housekeeping gene. One-way ANOVA with Tukey test, not significant (NS), * p-value < 0.03, ** p-value < 0.0001. Individual data point showing n=13 for NPC and n=4 for all other groups. **C** qPCR of immaturity (*Igfbp2*) genes. ΔΔct values normalized to NPC (100 ng/ml E/F) with *Rpl22-HA* used as a housekeeping gene. One-way ANOVA with Tukey test, not significant (NS), * p-value < 0.02, ** p-value < 0.0005. Individual data point showing n=13 for NPC and n=4 for all other groups. All graphs are mean±s.e.m.

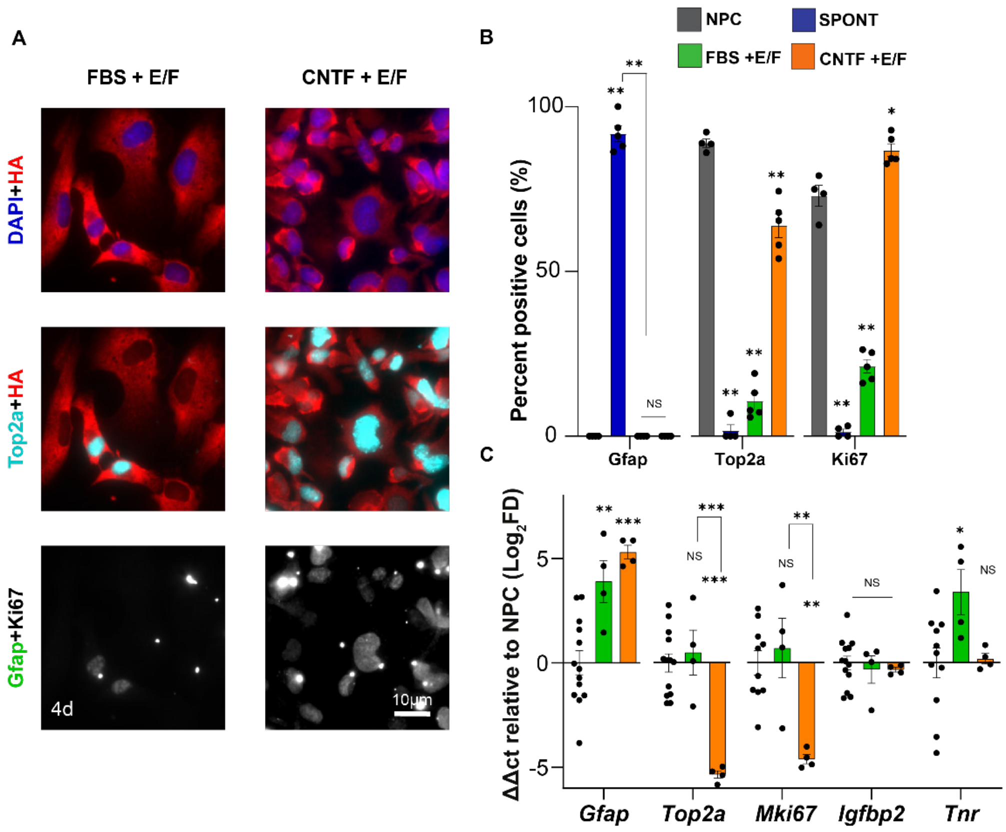


**Supplementary Figure 4.** Concurrent/simultaneous exposure of differentiation cytokines (CNTF, FBS) and EGF/FGF(E/F) do not yield astrocytes. **A** ICC staining of concurrent/simultaneously differentiated NPC for Ribo Tag (HA), nucleus (DAPI) astrocyte (Gfap), and proliferation markers (Top2a and Ki67), showing no astrocytes present, more proliferation in CNTF group and minimal proliferation in FBS group. **B** Quantification of percentage positive astrocytes (Gfap) and proliferative (Top2a, Ki67) cells from ICC. Graph shows mean± s.e.m, one-way ANOVA with Tukey test, not significant (NS), ** p-value < 0.03, *** p-value < 0.0001. Individual data point showing n=4 for all groups. **C** qPCR of astrocyte (*Gfap*), proliferative (*Top2a, Ki67*), immaturity (*Igfbp2*) and oligodendrocyte (*Tnr*) gene. ΔΔct values normalized to NPC (100 ng/ml E/F) with *Rpl22-HA* used as a housekeeping gene. One-way ANOVA with Tukey test, not significant (NS), * p-value 0.04, ** p-value < 0.006, *** p-value < 0.0003. Individual data point showing n=13 for NPC and n=4 for all other groups. All graphs are mean±s.e.m.

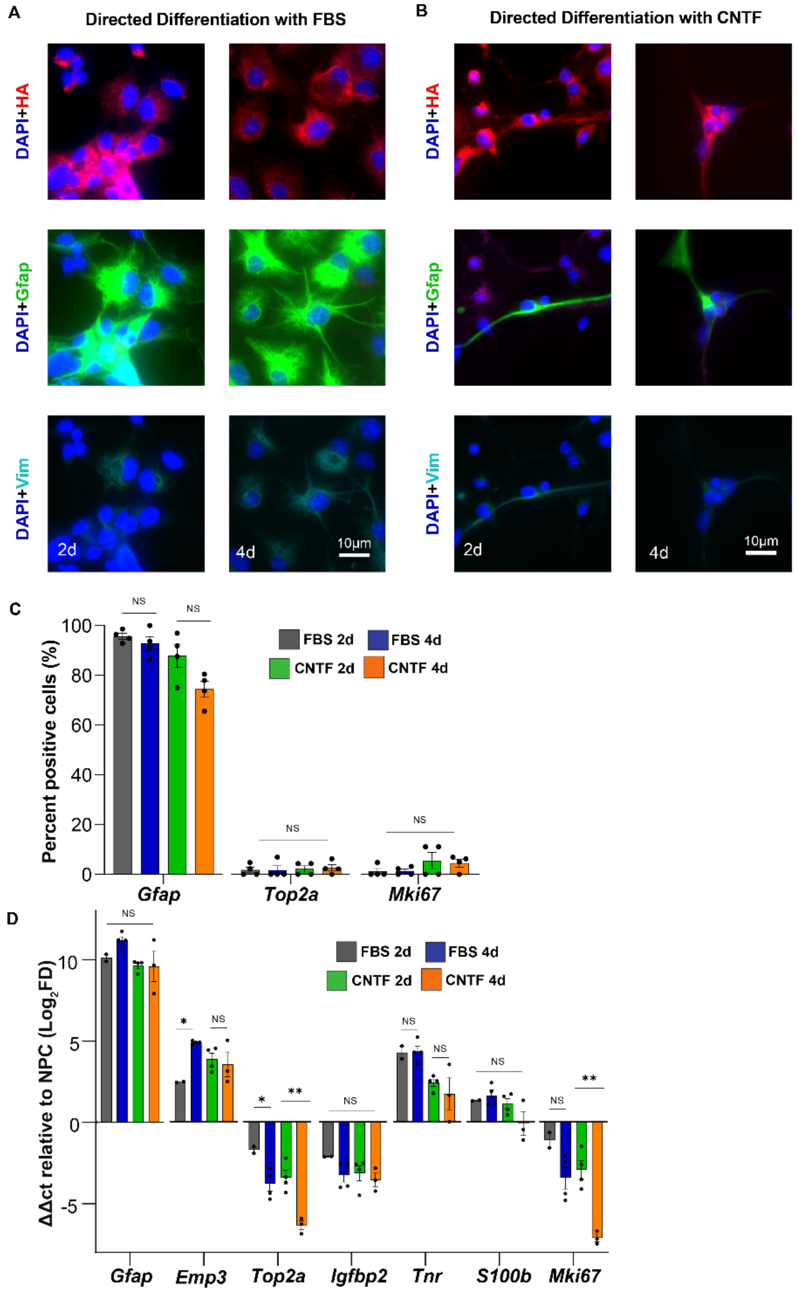


**Supplementary Figure 5.** Directed Differentiation occurs as early as 2 days with similar profiles to 4-day differentiation. **A&B** ICC staining of directed differentiated with FBS (A) and CNTF (B) NPC for Ribo Tag (HA), nucleus (DAPI), and astrocyte (Gfap, Vimentin), showing similar protein expressions. **C** Quantification of percentage positive astrocytes (Gfap) and proliferative (Top2a, Ki67) cells. Graph shows mean± s.e.m, one-way ANOVA with Tukey test, not significant (ns). Individual data point showing n=4 for all groups. **D** qPCR of astrocyte (*Gfap, Emp3, S100b),* proliferative (*Top2a, Ki67*), immaturity (*Igfbp2*) and oligodendrocyte (*Tnr*) gene. ΔΔct values normalized to NPC (100 ng/ml E/F) with *Rpl22-HA* used as a housekeeping gene. One-way ANOVA with Tukey test, not significant (NS), * p-value < 0.05, ** p-value < 0.004. Individual data point showing n=13 for NPC and n=4 for all other groups. **E** Percentage change in cell number of Immature astrocytes (ImA) at 2 days (Directed differentiation) and at 5 days (Induction of immaturity) showing that cells become proliferative according to cell counts when growth factors are reintroduced. Graph shows mean± s.e.m and individual data point showing n=4 for all groups. All graphs are mean±s.e.m.

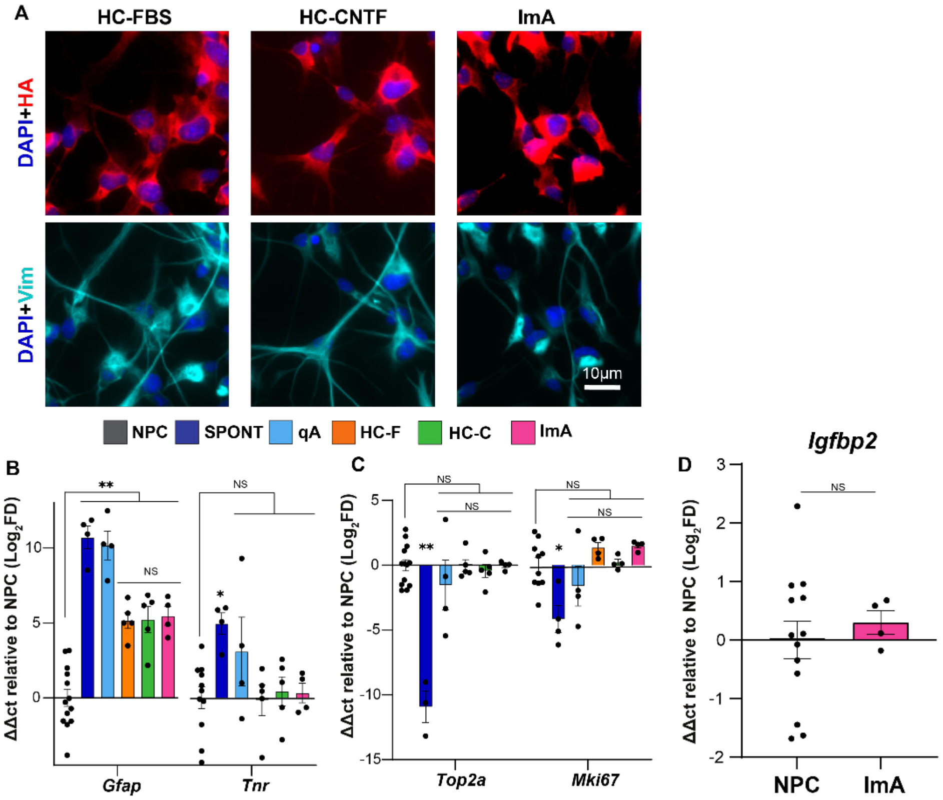


**Supplementary Figure 6**. Hysteretic conditioning (HC) of NPC generates immature astrocytes with high proliferation capacity. **A.** ICC detailed images after hysteretic conditioning of NPC with FBS (HC-FBS), CNTF (HC-CNTF) or CNT+FBS (ImA) showing the generation immature astrocytes staining for Vimentin. **B.** Hysteretic conditioning (HC) effects on *Gfap* and *Tnr* gene expression by qPCR. ΔΔct values normalized to NPC (100ng/ml E/F) with *Rpl22-HA* used as a housekeeping gene. One-way ANOVA with Tukey test, not significant (NS), * p-value < 0.03, ** p-value < 0.0001. Individual data point showing n=13 for NPC and n=4 for all other groups. All graphs are mean±s.e.m. **C.** Hysteretic conditioning (HC) effects on *Top2a* and *Mki67* gene expression by qPCR. ΔΔct values normalized to NPC (100 ng/ml E/F) with *Rpl22-HA* used as a housekeeping gene. One-way ANOVA with Tukey test, not significant (NS), * p-value < 0.007, ** p-value < 0.0001. Individual data point showing n=13 for NPC and n=4 for all other groups. All graphs are mean±s.e.m. **D** ImA express immaturity gene Igfbp2 at comparable levels to NPC. ΔΔct values normalized to NPC (100 ng/ml E/F) with *Rpl22-HA* used as a housekeeping gene. Student's t-test, not significant (NS). Individual data point showing n=13 for NPC and n=4 for ImA. All graphs are mean±s.e.m.

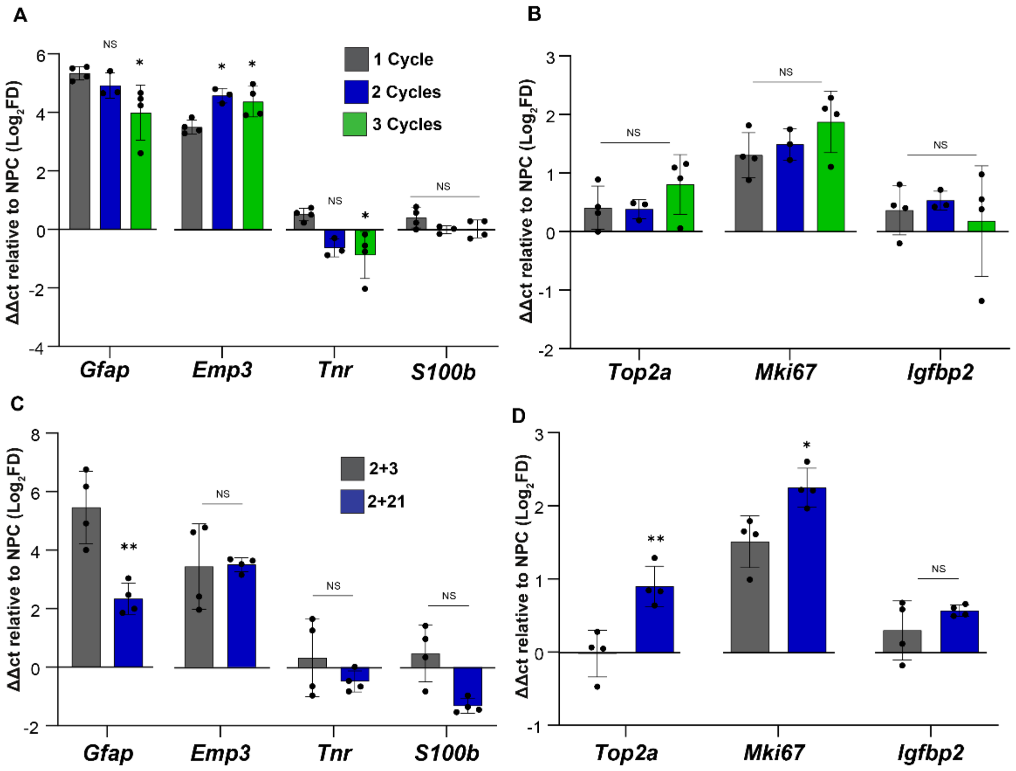


**Supplementary Figure 7**. Multiple cycles of Hysteresis yield similar profiles and induction of maturity occurs comparable at 3 days and 21 days. 1 Cycle; 2 days of directed differentiation, 2 days of immaturity induction. **A** qPCR of astrocytes (*Gfap, Emp3, S100b*) and oligodendrocytes (*Tnr*) genes. ΔΔct values normalized to NPC (100 ng/ml E/F) with *Rpl22-HA* used as a housekeeping gene. One-way ANOVA with Tukey test, not significant (NS), * p-value < 0.04. Individual data point showing n=4 for all. Showing similar profiles with multiple cycles. **B** qPCR of proliferation (*Top2a and Ki67*) and immaturity (*Igfbp2*) genes. ΔΔct values normalized to NPC (100 ng/ml E/F) with *Rpl22-HA* used as a housekeeping gene. One-way ANOVA with Tukey test, not significant (NS). Individual data point showing n=4 for all. Showing similar profiles with multiple cycles. **C** qPCR of astrocytes (*Gfap, Emp3, S100b*) and oligodendrocytes (*Tnr)* genes. ΔΔct values normalized to NPC (100ng/ml E/F) with *Rpl22-HA* used as a housekeeping gene. Student's t-test, not significant (NS), * p-value < 0.02, ** p-value < 0.004. Individual data point showing n=4 for all. **D** qPCR of proliferation (*Top2a and Ki67*) and immaturity (*Igfbp2*) genes. ΔΔct values normalized to NPC (100 ng/ml E/F) with *Rpl22-HA* used as a housekeeping gene. Student's t-test, not significant (NS), * p-value < 0.03, ** p-value < 0.005. Individual data point showing n=4 for all other groups. All graphs are mean±s.e.m

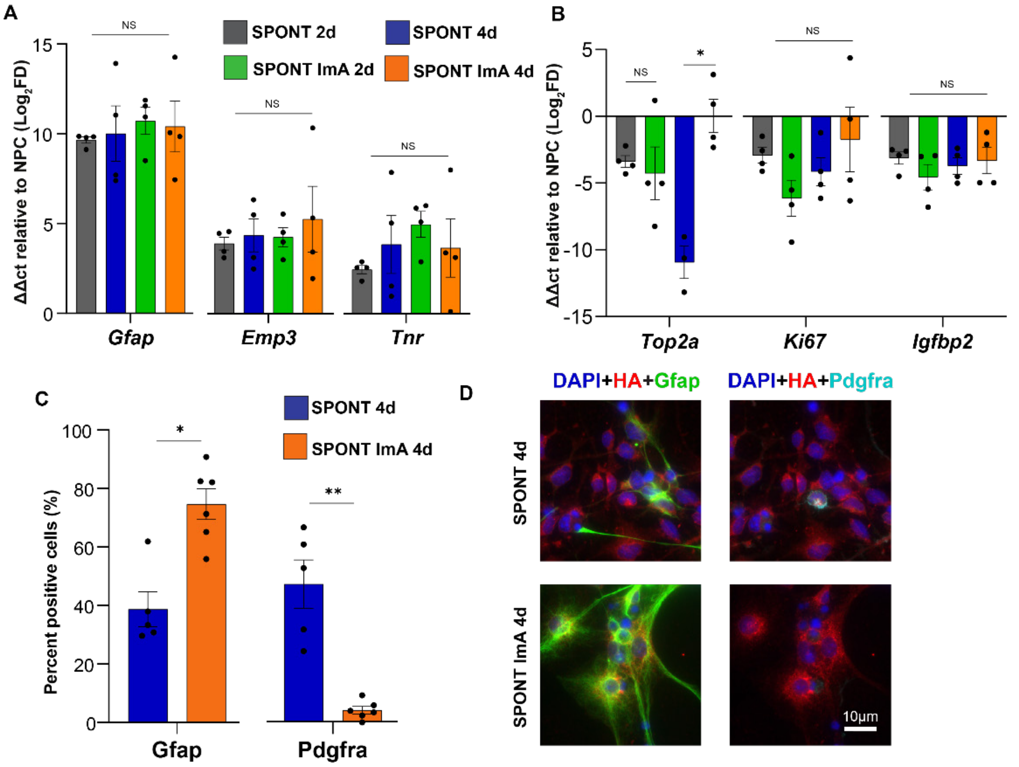


**Supplementary Figure 8**. Spontaneous differentiation of Immature astrocytes (ImA) has a higher retention of astrocyte character, lower oligodendrocyte yield and more proliferation capacity compared to spontaneously differentiated NPCs. **A** qPCR of astrocyte (*Gfap* and *Emp3*) and oligodendrocyte (*Tnr*) genes. ΔΔct values normalized to NPC (100 ng/ml E/F) with *Rpl22-HA* used as a housekeeping gene. One-way ANOVA with Tukey test, not significant (NS). Individual data point showing n=4 for all. **B** qPCR of proliferation (*Top2a* and *Ki67*) and immaturity (*Igfbp2*) genes. ΔΔct values normalized to NPC (100 ng/ml E/F) with *Rpl22-HA* used as a housekeeping gene. One-way ANOVA with Tukey test, not significant (NS), * p-value < 0.001. Individual data point showing n=4 for all. **C** Quantification of percentage positive astrocytes (*Gfap*) and oligodendrocytes (*Pdgfra*). Student's t-test, ** p-value < 0.002, *** p-value < 0.0003. Individual data point showing n=5 for all groups. **D** ICC Images of Spontaneously differentiated Immature astrocytes and NPCs showing glia marker, Gfap (astrocyte) and Pdgfra (oligodendrocyte), nuclei (DAPI) and Ribo Tag (HA). All graphs are mean±s.e.m.

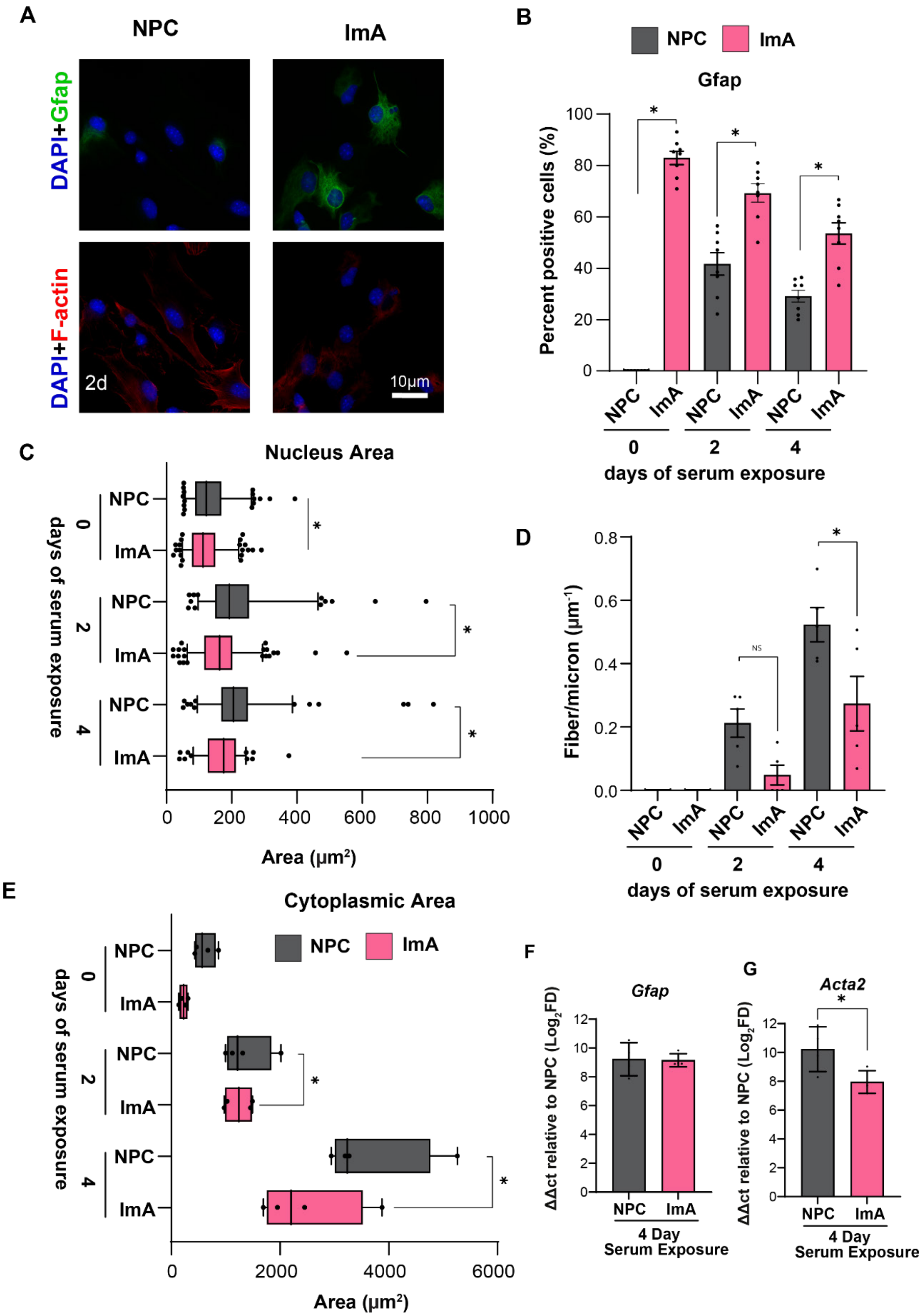
A
NPC
ImA
DAPI+Gfap
DAPI+F-actin
2d
10μm
B
NPC
ImA
Gfap
Percent positive cells (%)
100
80
60
40
20
0
NPC
ImA
0
2
4
days of serum exposure
C
Nucleus Area
NPC
ImA
days of serum exposure
0
2
4
0
200
400
600
800
1000
Area (μm²)
D
Fiber/micron (μm⁻¹)
0.8
0.6
0.4
0.2
0.0
NS
days of serum exposure
E
Cytoplasmic Area
0
2000
4000
6000
Area (μm²)
F
Gfap
ΔΔct relative to NPC (Log2FD)
4 Day
Serum Exposure
G
Acta2
12
10
8
6
4
2
0

**Supplementary Figure 9.** Gfap and Acta2/F-Actin expression, Actin Fiber alignment. Increased cytoplasmic and nuclear size as all pointers of EMT-like Transitions in NPC. Immature astrocytes are less likely to undergo these transitions. **A** ICC staining of NPC and ImA before and after serum exposure showing Ribo Tag (HA), Nucleus (DAPI) astrocytes (Gfap) and cytoskeletal filament (F-actin) markers showing reduced F-actin and retention of Gfap protein expression in ImA even at 2days of serum exposure. **B** Quantification of percentage positive astrocytes (Gfap) after 0, 2 and 4 days of serum exposure, showing that immature astrocytes predominantly retain their astrocyte character. Graph shows mean± s.e.m, one-way ANOVA with Tukey test, *** p-value < 0.0001. Individual data point showing n=7 for all groups. **C** Quantification of cell nucleus area at 0, 2 and 4 days of serum exposure percentage showing larger distribution of cell nucleus with time and greater distribution in NPC than ImA. Cell nucleus area calculated as area of DAPI intensity using FIJI Image J. Graph shows $25^{th}$ and 75th quartile, mean ± s.e.m, one-way ANOVA with Tukey test, *** p-value < 0.0001. Individual data point showing n=4 for all groups. **D** Quantification of F actin Fiber in cell cytoplasm at 0, 2 and 4 days of serum exposure showing increase in fiber density with time and less increase in ImA compared too NPC. Fiber per micron calculated as the number of peaks in a plot profile of a cell with F-actin ICC staining using FIJI Image J. Graph shows mean± s.e.m, one-way ANOVA with Tukey test, not significant (ns), * p-value < 0.01. Individual data point showing n=5 for all groups. **E** Quantification of cytoplasmic area at 0, 2 and 4 days of serum exposure percentage showing larger distribution of cell nucleus with time and greater distribution in NPC than ImA. Cytoplasmic area calculated as area of HA or F-actin intensity using FIJI Image J. Graph shows $25^{th}$ and 75th quartile ± s.e.m one-way ANOVA with Tukey test, *** p-value < 0.0001. Individual data point showing n=5 for all groups. **F** qPCR of astrocyte (*Gfap*) gene. ΔΔct values normalized to NPC (100ng/ml E/F) with *Rpl22-HA* used as a housekeeping gene. Graph shows mean± s.e.m, student's t-test. Individual data point showing n=4 for all groups. **G** qPCR of smooth muscle actin (*Acta2*) gene. ΔΔct values normalized to NPC (100 ng/ml E/F) with *Rpl22-HA* used as a housekeeping gene. Student's t-test, not significant (NS), * p-value < 0.04. Individual data point showing n=4 for all groups. All graphs are mean±s.e.m.

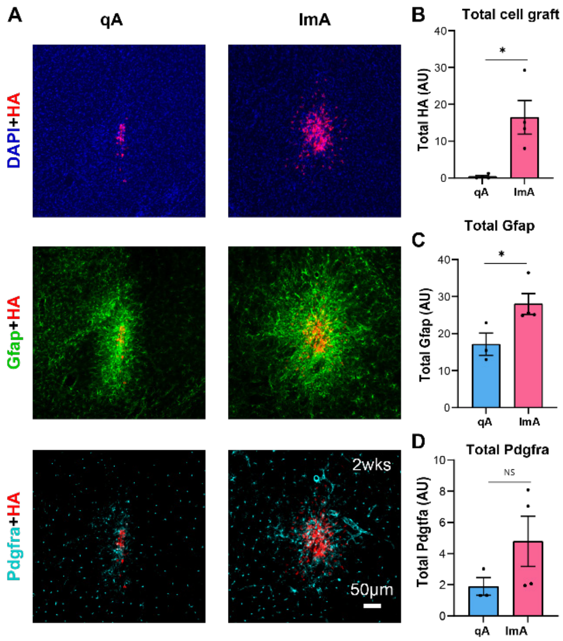


**Supplementary Figure 10**. Cell grafts in healthy mouse striatum. **A** IHC images at 2 weeks post cell graft showing grafted cells (HA), astrocytes (Gfap), oligodendrocytes (Pdgfra) and nuclei (DAPI). **B** Total cell graft quantified by HA Intensity. Graph shows mean± s.e.m, student's t-test, * p-value < 0.005. **C** Total intensity from cell injection site. Graph shows mean± s.e.m , student's t-test, * p-value < 0.005. **D** Total pdgfra intensity at injection site. Graph shows mean± s.e.m , student's t-test, not significant (NS).

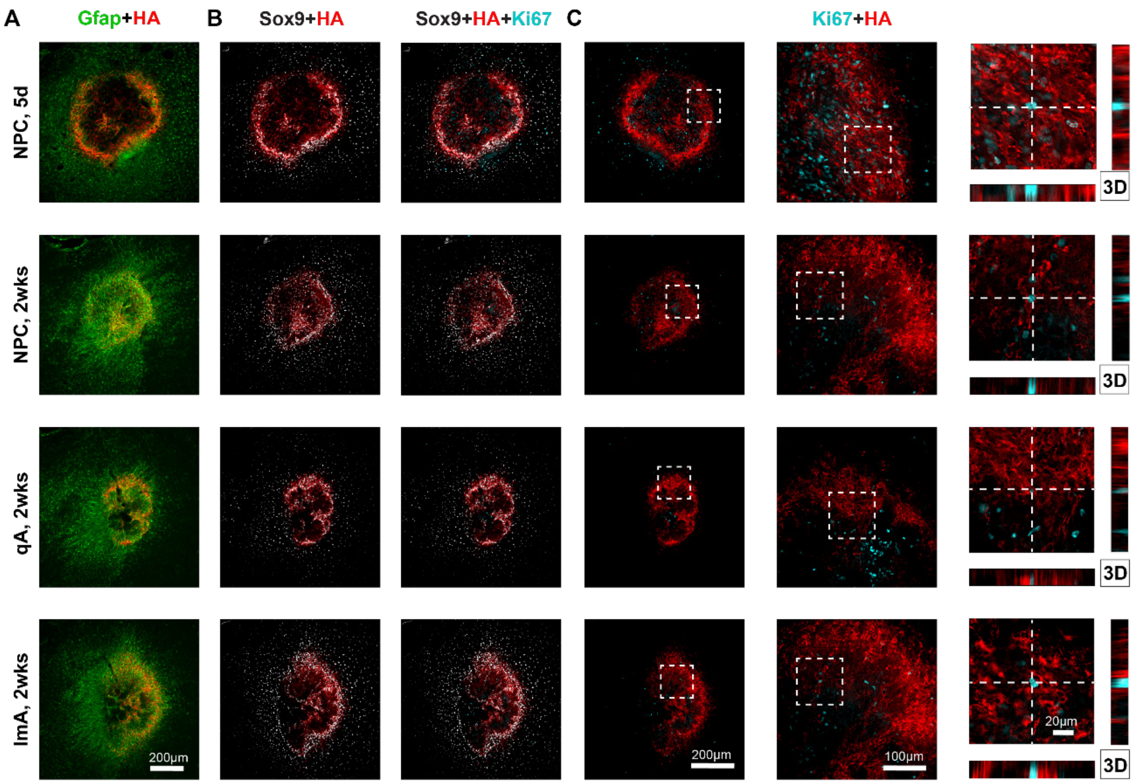


**Supplementary Figure 11**. Grafted cells do not proliferate at lesion site. **A&B** IHC images at 5d and 2 weeks post stroke showing grafted cells (HA), astrocytes (Gfap, Sox9) and proliferative cells (Ki67). **C** High magnification 3D IHC images showing no colocalization between NPC grafts (HA) and proliferation marker (Ki67) and at 5 days and 2 weeks for all graft types.

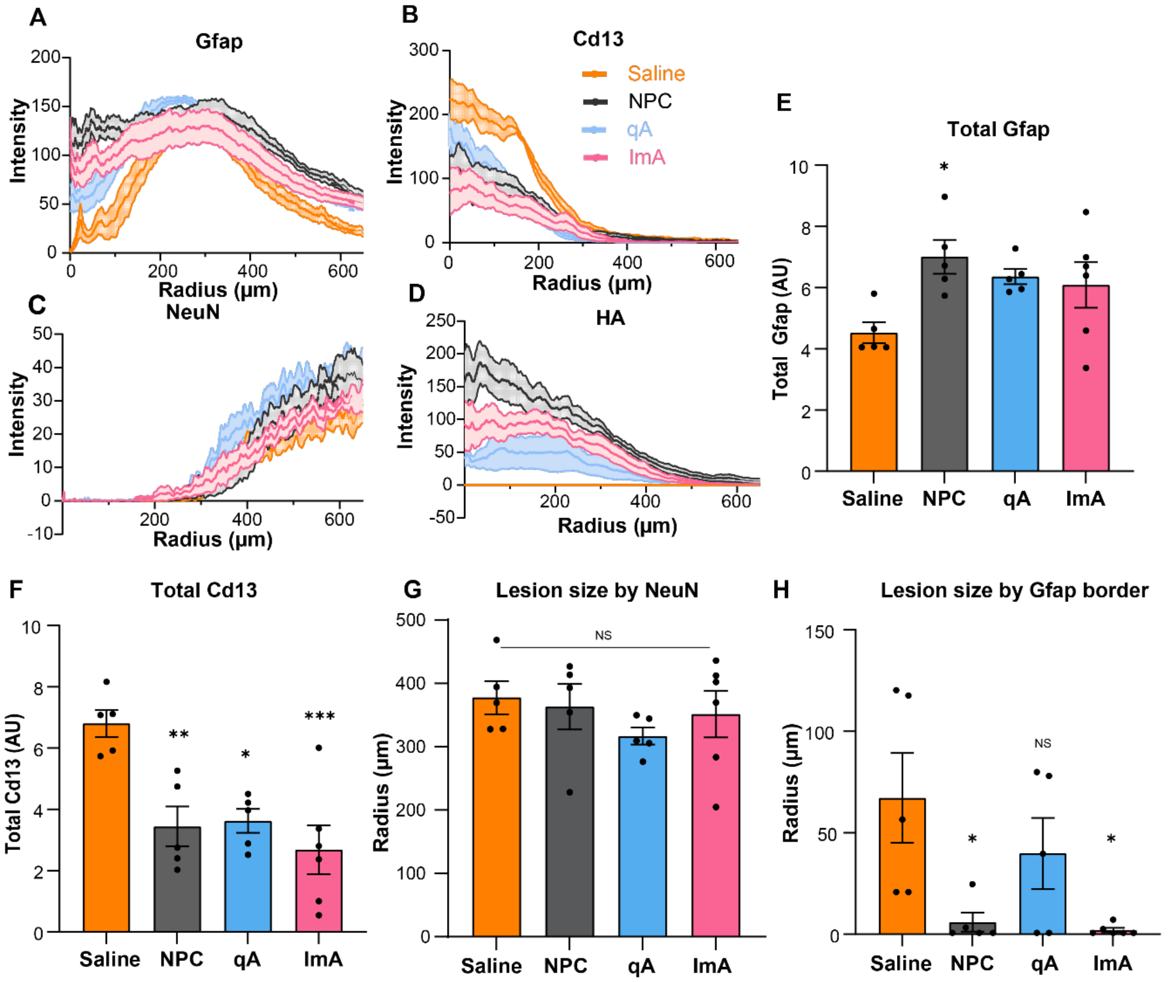


**Supplementary Figure 12**. Traces from center of lesion in ischemic stroke, cell grafts impact host immune response but does not affect lesion size **A** Quantification of Cd13 intensity radially from center the of the lesion, plot shows mean±s.e.m. **B** Quantification of Gfap intensity radially from center the of the lesion, plot shows mean±s.e.m. **C** Quantification of HA intensity radially from center the of the lesion, plot shows mean±s.e.m. **D** Quantification of NeuN intensity radially from center the of the lesion, plot shows mean±s.e.m. **E** Quantification Gfap expression at stroke lesions using cumulative Gfap intensity in ischemic stroke. Total Gfap intensity calculated as area under the curve (AUC) up to 350 µm radially into the tissue per animal. Graph shows mean±s.e.m, one-way ANOVA with Tukey test, * p-value < 0.05. Individual data point showing n=5 for saline, NPC and qA and n=6 for ImA. **F** Quantification Cd13 expression at stroke lesions using cumulative Cd13 intensity in ischemic stroke. Total Cd13 intensity calculated as area under the curve (AUC) up to 350 µm radially into the tissue per animal. Graph shows mean±s.e.m, one-way ANOVA with Tukey test, * p-value < 0.05, ** p-value < 0.05, *** p-value < 0.05. Individual data point showing n=5 for saline, NPC and qA and n=6 for ImA. **G** Lesion size quantified by NeuN intensity showing that neurons are not present in center of lesion up to about 350 µm radially. Graph shows mean±s.e.m, one-way ANOVA with Tukey test, not significant (NS). **H** Lesion size quantified by Gfap intensity showing that NPC and ImA fill lesion core with Gfap-positive cells. Graph shows mean±s.e.m, one-way ANOVA with Tukey test, * p-value < 0.05.

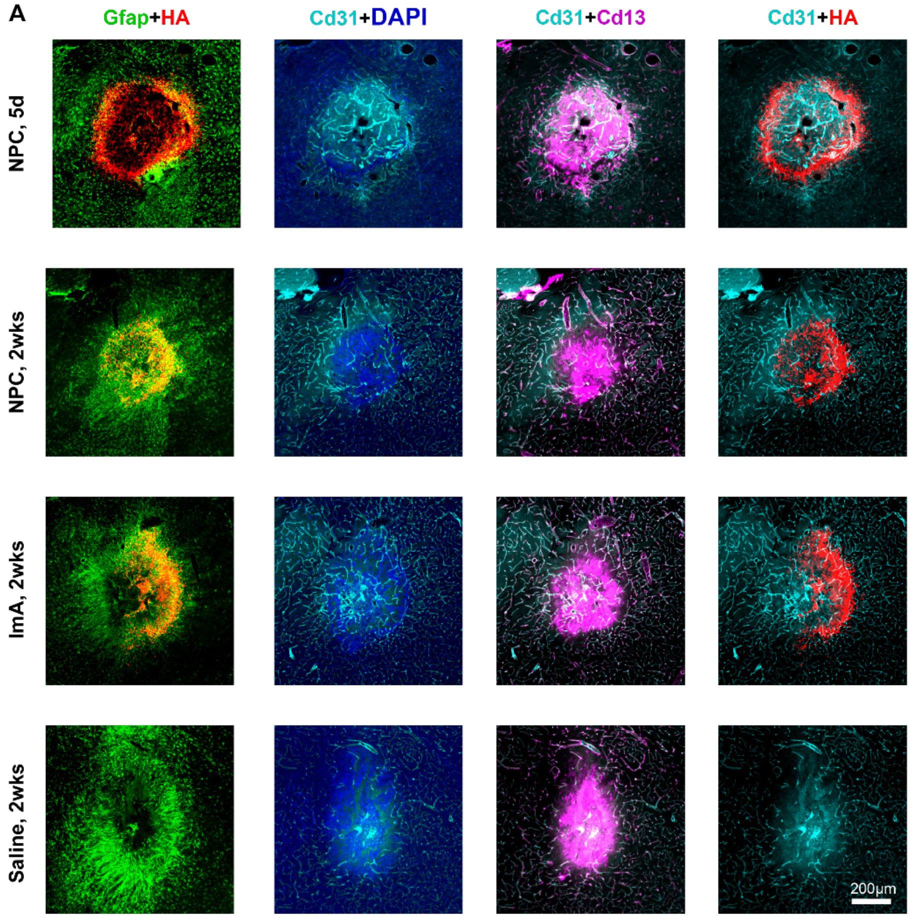


**Supplementary Figure 13**. Cell grafts in ischemic strokes encourage angiogenesis observed at 2 weeks. **A** IHC images at 2 weeks post stroke showing grafted cells (HA), astrocytes (Gfap), nuclei (DAPI), myeloid lineage cells (Cd13) and platelet endothelial cell adhesion molecule (Cd31). Showing Cd31 lined vasculature in ImA and NPC grafted lesions at 2 weeks with no defined structure of vessel in saline at 2 weeks.

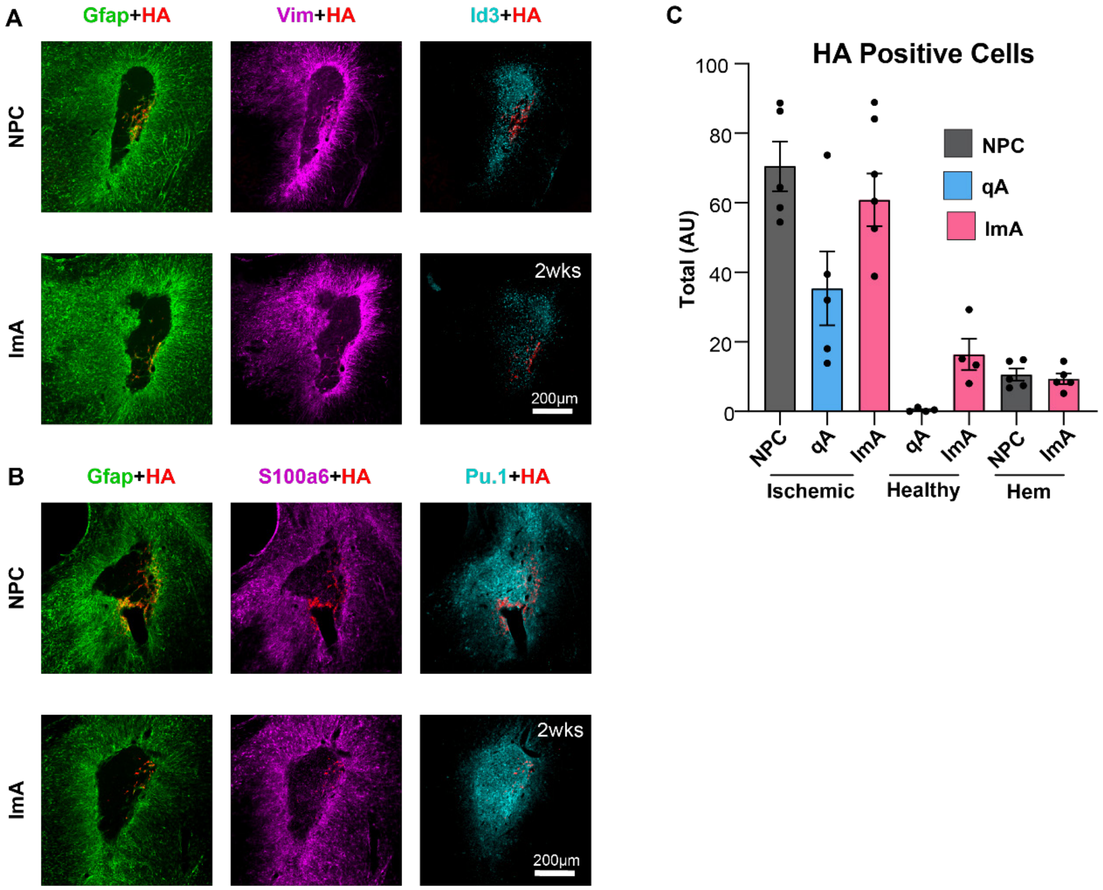


**Supplementary Figure 14**. Cell grafts in hemorrhagic stroke express border forming astrocyte markers. **A** IHC images at 2 weeks post stroke showing grafted cells (HA), astrocytes (Gfap, Vimentin) and border forming astrocytes (Id3). **B** IHC images at 2 weeks post stroke showing grafted cells (HA), astrocytes (Gfap) and border forming astrocytes (S100a6 and Pu.1). **C** Quantification of surviving grafts at ischemic, hemorrhagic stroke lesions and healthy mouse striatum with NPC, qA and ImA, using cumulative HA intensity in ischemic stroke derived from cumulative intensity measured radially from the center of lesion and normalized to total integrated density. Total HA intensity calculated as area under the curve (AUC) up to 650 μm radially into the tissue per animal. Graph shows mean±s.e.m, individual data point showing n=5 for saline, NPC and qA and n=6 for ImA.

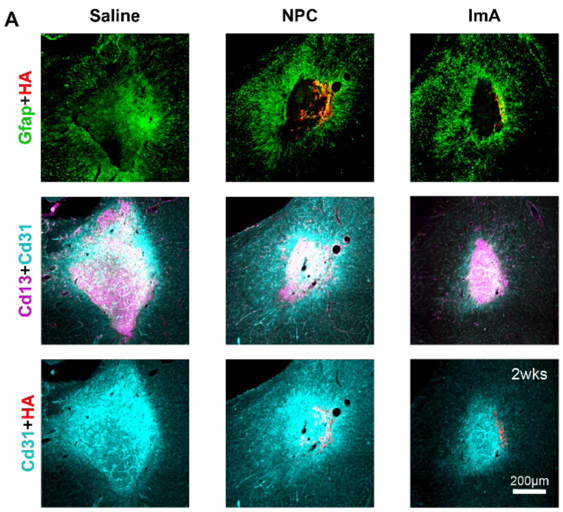


**Supplementary Figure 15**. Grafts have no effect on angiogenic activity at center of lesion. **A** IHC images at 2 weeks post stroke showing grafted cells (HA), astrocytes (Gfap), platelet endothelial cell adhesion molecule (Cd31) and myeloid lineage cells (Cd13). Cd31 showing diffuse staining in all conditions.

| | | |
|---|---|---|
| ***RiboTag (HA)*** | forward | GAACAACCTCCGAGACTGGC |
| | reverse | CATAGTCCGGGACGTCATAGGG |
| ***Gfap*** | forward | CGGAGACGCATCACCTCTG |
| | reverse | AGGGAGTGGAGGAGTCATTCG |
| ***S100b*** | forward | TGGTTGCCCTCATTGATGTCT |
| | reverse | CCCATCCCCATCTTCGTCC |
| ***Top2a*** | forward | CAACTGGAACATATACTGCTCCG |
| | reverse | GGGTCCCTTTGTTTGTTATCAGC |
| ***Igfbp2*** | forward | CAGACGCTACGCTGCTATCC |
| | reverse | CCCTCAGAGTGGTCGTCATCA |
| ***Acta2*** | forward | GTCCCAGACATCAGGGAGTAA |
| | reverse | TCGGATACTTCAGCGTCAGGA |
| ***Mki67*** | forward | ATCATTGACCGCTCCTTTAGGT |
| | reverse | GCTCGCCTTGATGGTTCCT |
| ***Tnr*** | forward | GGCTGGAGGTGACTACAGAAA |
| | reverse | GAAGACCATAGGCTGTTCCTTG |
| ***Emp3*** | forward | TGGTGCTGTCTCTCATCCTCT |
| | reverse | CGAAGCAGTAACCGAAGCTG |

**Supplementary Table 1**: List of primers used for qPCR in this study.
